# Evaluating the Analytical Performance of Direct-to-Consumer Gut Microbiome Testing Services

**DOI:** 10.1101/2024.06.05.596628

**Authors:** Stephanie L. Servetas, Diane Hoffmann, Jacques Ravel, Scott A. Jackson

## Abstract

Consumer interest in personal microbiome health has given rise to numerous direct-to-consumer (DTC) microbiome testing services despite questions regarding their analytical and clinical validity, and consumer safety. These tests straddle the line between more strictly regulated medical devices and minimally regulated general health and wellness products; a distinction that may not be readily apparent to consumers. To assess the current state of the industry, we evaluated the performance of seven commercial DTC gut microbiome testing services using a standardized NIST – developed human fecal material. Our results reveal major discrepancies, both within and across the different service providers. Significantly, we found variability between providers was on the same scale as biological variability between different donors. We attribute the observed differences to methodological variability and lack of sufficient quality control. Additionally, we highlight that analytical performance is a prerequisite for making sound clinical recommendations. Our results demonstrate the need for standards to ensure analytical validity and regulatory oversight to ensure patient safety.

## Introduction

Over the last decade, the human gut microbiome has been linked to a plurality of health and disease conditions including mental health, cancer, and obesity to name a few [1-3]. Despite extensive research, there are currently no regulatory-approved clinical microbiome diagnostic tests. Concurrently, direct-to-consumer (DTC) personal wellness services are becoming increasingly accessible despite challenges regarding their technical and clinical validity [4-6]. As our current understanding of the microbiome structure-function relationship is still limited, our ability to make confident diagnostic or prognostic predictions based on an individual’s microbiome is limited. The French Society of Microbiology, through a publication in Le Monde, has advised against microbiome testing, primarily due to insufficient knowledge and the personalized nature of defining a “healthy” microbiome [7].

Despite these challenges, associations between the gut microbiome and human health are often prominently featured by media outlets, leading many consumers to seek a better understanding of their own gut health. While citizen science projects like The Microsetta Initiative have offered consumers basic information about their gut microbiomes, there remains a demand for more actionable information about the status and implications of personal gut health [8, 9]. In response, numerous commercial DTC microbiome testing services have emerged, providing consumers with profiles of their gut microbiome. Many of these companies extend their services beyond microbiome profiling by classifying microbes as pathogens or beneficial, comparing the profile of a consumer’s microbiome against a comparative population, providing an index of gut microbiome health, or offering recommendations for lifestyle changes, dietary changes and/or dietary supplements.

While these recommendations are often limited to dietary interventions, regulatory concerns regarding consumer protection have been raised [10]. Notably, there has been a technological shift from end-to-end diagnostic testing, traditionally confined to clinical settings and conducted by trained medical professionals, to DTC testing services like 23andMe. These services have introduced home testing kits that allow consumers (patients) to collect their own samples, ship them to a laboratory, and access the results without consulting a clinician. A primary concern is that these at-home tests do not undergo the same rigorous oversight in validating analytical performance as traditional medical diagnostic tests. Such validation is crucial to assure clinicians, patients, and regulators of the reliability and actionability of test results. This lack of validation and the implications may not be apparent to the consumer.

In the age of digital publishing, numerous citizen scientists, reporters, and microbiome researchers have submitted samples to different companies and published on their results [11-13]. While these case studies are often intriguing and provide some anecdotal insight, to our knowledge, a rigorous evaluation of the analytical performance has not yet been conducted. The National Institute of Standards and Technology (NIST) has developed a suite of candidate human fecal standards, confirmed to be homogeneous and stable via rigorous multi’omic analyses [14, 15]. These materials have been designed to enable stakeholders to evaluate the impact of methodological variability on their microbiome measurements, both qualitatively and quantitatively. The fecal standards, being homogenous and stable, are ideally suited to assess the precision (reproducibility) of measurement workflows within and across laboratories.

In this study, we describe our approach to evaluating the performance of seven DTC microbiome testing services using a NIST-developed human fecal standard. Each service employed an NGS-based analysis workflow (16S rRNA gene amplicon sequencing or whole metagenome shotgun sequencing (WMS)) and provided a detailed report of analysis. We assessed the reproducibility of the results, both within and across these testing services.

## Results

### Experimental Design Strategy

As part of its mission to promote U.S. innovation and industrial competitiveness, NIST has been working with stakeholders in industry, academia, and other government agencies to develop standards to support the advancement and commercial translation of microbiome science. To this end, we wished to assess the current state-of-the-art with respect to DTC gut microbiome testing services. To achieve this, we deployed a NIST-developed human gut microbiome (human fecal) standard to assess the precision/reproducibility of seven DTC gut microbiome testing companies. For each of the seven companies, three tests/kits were ordered via the company’s website. Upon receipt of the (3 x 7 = 21) sample collection kits, we used a homogenized pool of human fecal material to serve as the test sample material for each collection kit. Collection procedures were followed according to instructions provided by each company (Table 1). Samples were placed into a company-provided shipping container and returned to the company for analysis. Results, in the form of company-specific reports, were received anywhere from 2-8 weeks following shipment of the sample. Taxonomic profiles were manually extracted from these reports and used for all subsequent analyses. It is also noteworthy that the companies were not informed of the ongoing assessment until after all the samples had been processed and final reports were received by NIST.

**Table 1.**
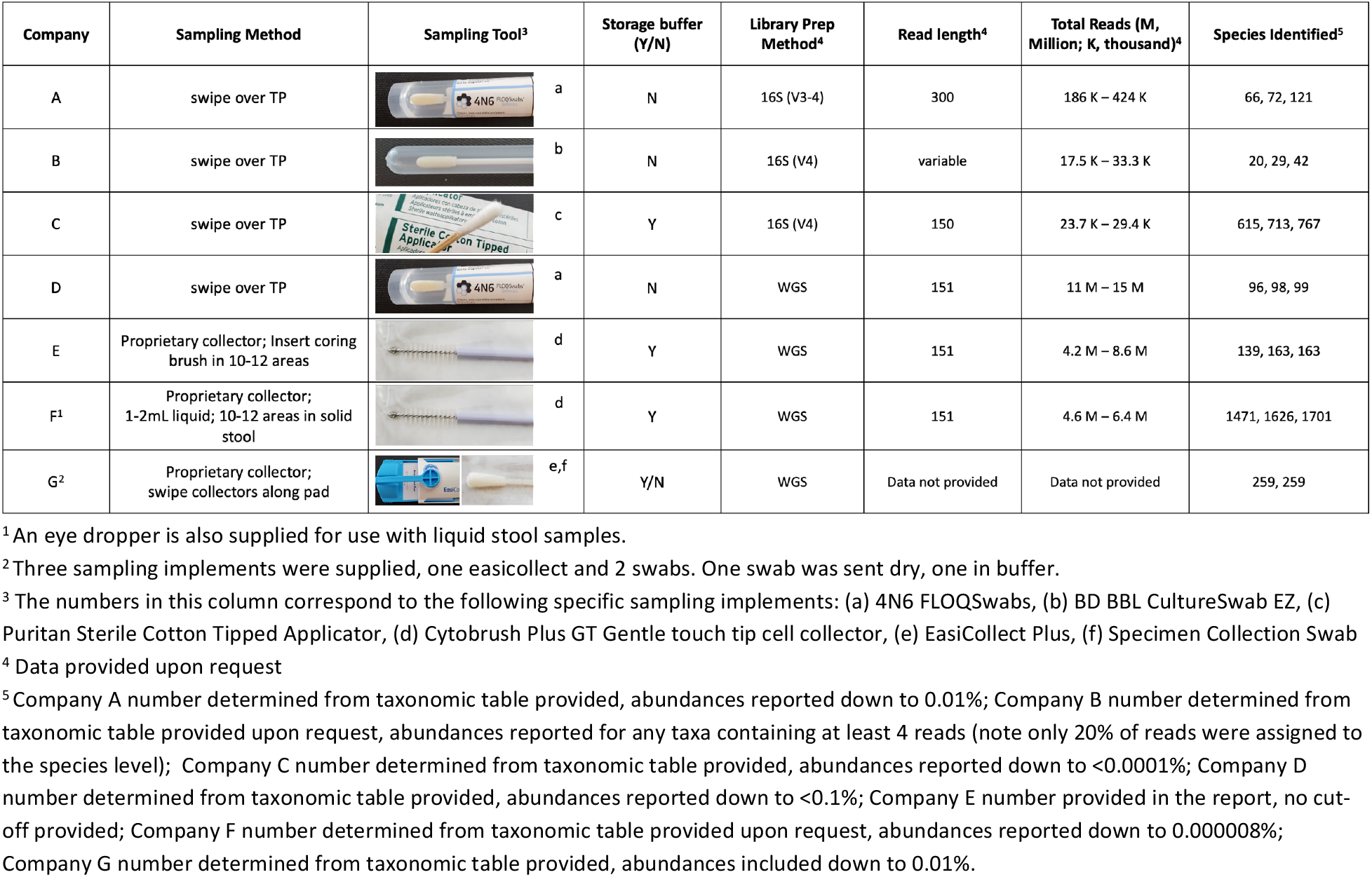
Gut Microbiome Testing Methodological Parameters.

| Company | Sampling Method | Sampling Tool<sup>3</sup> | Storage buffer (Y/N) | Library Prep Method<sup>4</sup> | Read length<sup>4</sup> | Total Reads (M, Million; K, thousand)<sup>4</sup> | Species Identified<sup>5</sup> |
| --- | --- | --- | --- | --- | --- | --- | --- |
| A | swipe over TP | a | N | 16S (V3-4) | 300 | 186 K – 424 K | 66, 72, 121 |
| B | swipe over TP | b | N | 16S (V4) | variable | 17.5 K – 33.3 K | 20, 29, 42 |
| C | swipe over TP | c | Y | 16S (V4) | 150 | 23.7 K – 29.4 K | 615, 713, 767 |
| D | swipe over TP | a | N | WGS | 151 | 11 M – 15 M | 96, 98, 99 |
| E | Proprietary collector; Insert coring brush in 10-12 areas | d | Y | WGS | 151 | 4.2 M – 8.6 M | 139, 163, 163 |
| F<sup>1</sup> | Proprietary collector; 1-2mL liquid; 10-12 areas in solid stool | d | Y | WGS | 151 | 4.6 M – 6.4 M | 1471, 1626, 1701 |
| G<sup>2</sup> | Proprietary collector; swipe collectors along pad | e,f | Y/N | WGS | Data not provided | Data not provided | 259, 259 |

<sup>1</sup> An eye dropper is also supplied for use with liquid stool samples.
<sup>2</sup> Three sampling implements were supplied, one easicollect and 2 swabs. One swab was sent dry, one in buffer.
<sup>3</sup> The numbers in this column correspond to the following specific sampling implements: (a) 4N6 FLOQSwabs, (b) BD BBL CultureSwab EZ, (c) Puritan Sterile Cotton Tipped Applicator, (d) Cytobrush Plus GT Gentle touch tip cell collector, (e) EasiCollect Plus, (f) Specimen Collection Swab
<sup>4</sup> Data provided upon request
<sup>5</sup> Company A number determined from taxonomic table provided, abundances reported down to 0.01%; Company B number determined from taxonomic table provided upon request, abundances reported for any taxa containing at least 4 reads (note only 20% of reads were assigned to the species level); Company C number determined from taxonomic table provided, abundances reported down to <0.0001%; Company D number determined from taxonomic table provided, abundances reported down to <0.1%; Company E number provided in the report, no cut-off provided; Company F number determined from taxonomic table provided upon request, abundances reported down to 0.000008%; Company G number determined from taxonomic table provided, abundances included down to 0.01%.

### Methodological parameters varied between companies

The process of translating the composition of stool sample into a microbiome report is multifaceted. As there are currently no universally accepted best practices for these methods, each provider likely employs a unique and often proprietary workflow. Microbiome measurements are impacted by a vast number of methodological variables including 1) sample collection, storage, and shipping methods, 2) nucleic acid extraction techniques, 3) NGS-library preparation, 4) sequencing technology, and 5) bioinformatic analyses. Bias can be introduced at every step, and even minor changes in methodology can lead to significant differences in results. In this study we had limited knowledge of the workflows that were employed by each company; however, we were able to infer key differences in both sample collection and sequencing methodologies.

Table 1 shows a comparison between companies regarding some methodological variables likely to impact the result (Table 1). Notably, there was considerable variability in methods for sample collection, including both the sample type (whole bowel movement vs. used toilet paper) and the sampling implement. Furthermore, some companies instructed users to add the sampling implement to a buffer, or to resuspend the sample in a buffer and discard the implement, while others directed users to ship the implement back ‘neat’, enclosed in a secondary sterile container. None of the shipping packages included ice, so all samples were shipped under ambient conditions (the study was conducted in the spring). The exact transit times between shipping and receipt of samples were not reported by all companies. Typically, notification of sample receipt or processing was sent 5-7 days after shipping.

Regarding sequencing methodologies, some companies used marker gene amplicon sequencing while others relied on WMS. Among those using amplicon sequencing, one utilized 16S rRNA gene amplicon sequencing of the V3-4 region and the other two used 16S rRNA gene amplicon sequencing of the V4 region; additionally, one company included sequencing of internal transcribed spacer (ITS) amplicon for fungal identification. The read length and depth of sequencing, which can impact the specificity and sensitivity of taxonomic identification, varied by company. Although a wide range of read depths was reported, the variability within a single company was generally narrower (Table 1). Read depth ranged from < 2 x 10^4^ to > 1 x 10^7^. Interestingly, higher read depths did not always correlate with a greater number of species identified. For example, the three replicates from company C each reported over 600 species with read depths between 2 x 10^4^ and 3 x 10^4^. In contrast, company D reported 95, 98, 99 species for their three replicates, each with read depths exceeding 1 x 10^7^ reads. Although most details of read processing were not disclosed, we noted different cutoffs (ranging from 0.1% to 8 x 10^-6^ %) for reporting taxonomic assignment based on the taxonomic tables provided.

### Taxonomic Profiles Varied Across Companies

The process of generating the taxonomic tables from the data provided by each company required manual curation, as the reports came in varied formats (e.g., PDF, html). Additionally, a complete taxonomic profile was not part of Company A’s report but was obtained after sending a request to their help center. For comparison, NIST also performed metagenomic analyses on aliquots of the same material that was shared with the DTC companies. In addition, taxonomic data from the other 7 unique donors that contributed to the candidate fecal reference material development were included to highlight the differences associated with biological variability (donor) *vs*. technical (methodological) variability. Although species-levels taxa were reported by each DTC company, all subsequent analyses at NIST were conducted using genus-level taxa assignments. A comparison of alpha diversity revealed a range of diversity values with no clear patterns based on read depth or sequencing method (Figure 1A-C, respectively). Bray-Curtis dissimilarity was calculated to compare all replicates from each of the seven companies and NIST (Figure 1D). Along with the alpha-diversity plots, the ordination plot can indicate inter-lab reproducibility. Narrower box plots (Figure 1A-C) and tighter distribution on the ordination plot (Figure 1D) suggest higher reproducibility for a company (*e*.*g*., Company F), while Company A showed poor reproducibility. Moreover, there was no distinctive clustering based on whether WMS or amplicon sequencing was used (WMS shown with solid lined, and 16S rRNA gene amplicon sequencing with dashed lined ellipses in Figure 1D). One question raised was how the diversity observed for a single sample analyzed by multiple companies compared to biological diversity from different donors analyzed by a single workflow. A second Bray-Curtis dissimilarity index was calculated that included data from seven additional donors processed with the NIST workflow. Surprisingly, the diversity observed for samples from a single donor analyzed by different companies (asterisk in Figure 1E, blue ellipses) appeared to be greater than or equal to that of biologically distinct samples (Figure 1E, other shapes, yellow ellipse). This demonstrates that methodological variability can be greater than biological variability.

**Figure 1.**
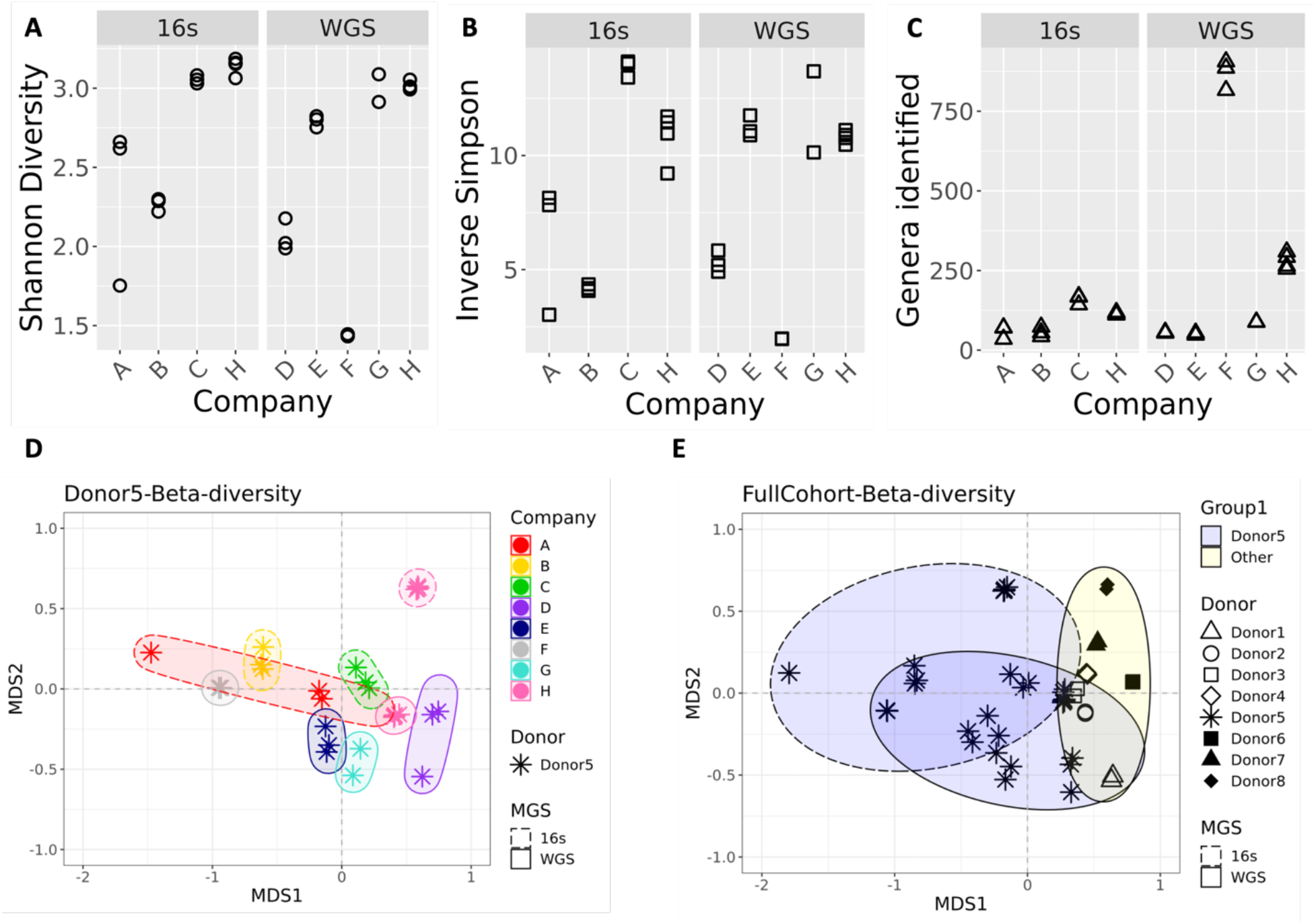
Diversity Metrics for Genera identified by DTC companies and by NIST. Plots show alpha-diversity calculated using **(A)** Shannon **(B)** Inverse Simpson indices and **(C)** for the total genera identified. Three replicates are plotted for companies A, B, C, D, E, and F; company G has 2 replicates and company H has 4. **(D)** Ordination plot showing beta-diversity based on genera for different companies processing samples from a single donor. Colors indicated different Companies; ellipses are drawn to further identify samples processed with the same Company. Ellipses with dashed lines indicates 16S amplicon sequencing was used; solid lines on the ellipses indicate WMS. **(E)** Ordination plot showing beta-diversity based on genera for a single donor processed by different companies (asterisk) and different donors processed by a single workflow (other symbols). The yellow ellipses indicate the cluster of other donors processed with a single workflow; blue ellipses indicate samples from a single donor. Similar to plot D, dashed lined ellipses denote 16S, solid denote WMS

In addition to comparing diversity metrics, the taxonomic composition was compared on a genus-by-genus basis to identify the genera common to all analyses. For this analysis, any taxa that was not given a specific genus designation was renamed unspecified and these assignments were then grouped with any unidentified taxa that were noted in the report. While some companies provided information that allowed us to determine the percentage of unidentified reads, this was not uniform across companies. In fact, the NIST analysis was based on Bracken, and in this workflow unidentified reads are dropped prior to calculating relative abundance. Therefore, while we were able to show this data for some companies, absence of the unspecified category in Figure 2 does not indicate 100% of the taxa were identified but could indicate that the unidentified reads were dropped earlier in the process, prior to calculating relative abundance values for known taxa. Despite vast differences in read depths, it was hypothesized that the most common genera would be present in all samples, and less abundant genera would only be seen in samples with higher coverage. The number of identified genera ranged from 34 to 906, with a relatively narrow range for each single company (Figure 1C). From all 9 analyses (7 companies and NIST WMS and 16S rRNA gene amplicon sequencing), a total of 1208 unique taxa were identified. Only 3 genera were found in all samples when comparing all taxonomic profiles (data not shown). A single replicate from Company A had a distinctive taxonomic profile (Figure 3B, and observed in Figure 1D as the single red asterisk on the left), which when removed from the analysis, increased the number of common genera to 17. This accounted for less than 2 % of the total genera identified, or between 2 % and 43 % for any given sample. In most cases, these common genera constituted a majority of the genera (> 50 % of the abundance for companies A, C, D, G, and both NIST analyses), or the majority of genus level specified reads (companies B and F). Additionally, the conservation of identified genera between replicates varied, from as low as 6 % for Company A to as high as 98 %. Excluding Company A, the genera conserved between replicates from the same company accounted for at least 95% of the identified sample composition. This indicates that most variability between replicates from a single company occurs among taxa present at low abundance and highlights the importance of empirically determined read cut-offs for taxa inclusion in a report.

**Figure 2.**
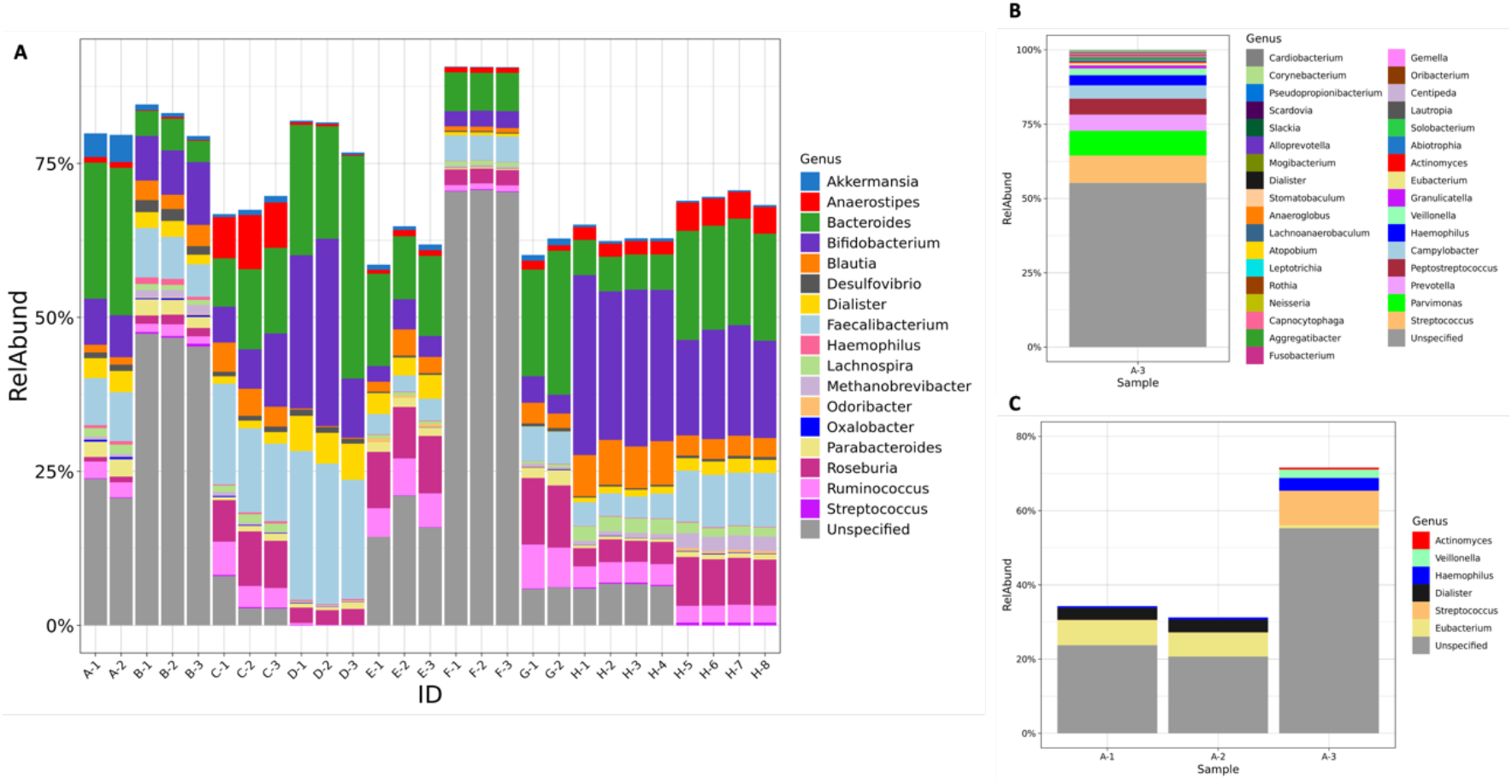
Common genera identified across testing companies. (A) The relative abundance of genera that were identified in all but one test. Each company is indicated by a letter, with replicate samples indicated by the number after the dash. (B) Relative abundance data for the one test sample (A-3) was excluded from the overall analysis. All genera that were identified are included in the plot, (C) Relative abundance of genera found in common across replicates run by company A. The grey portion in all bars represents taxa that were not given a Genus level identification; this includes both taxa designated at a higher taxonomic level or that were listed as unidentified.

**Figure 3.**
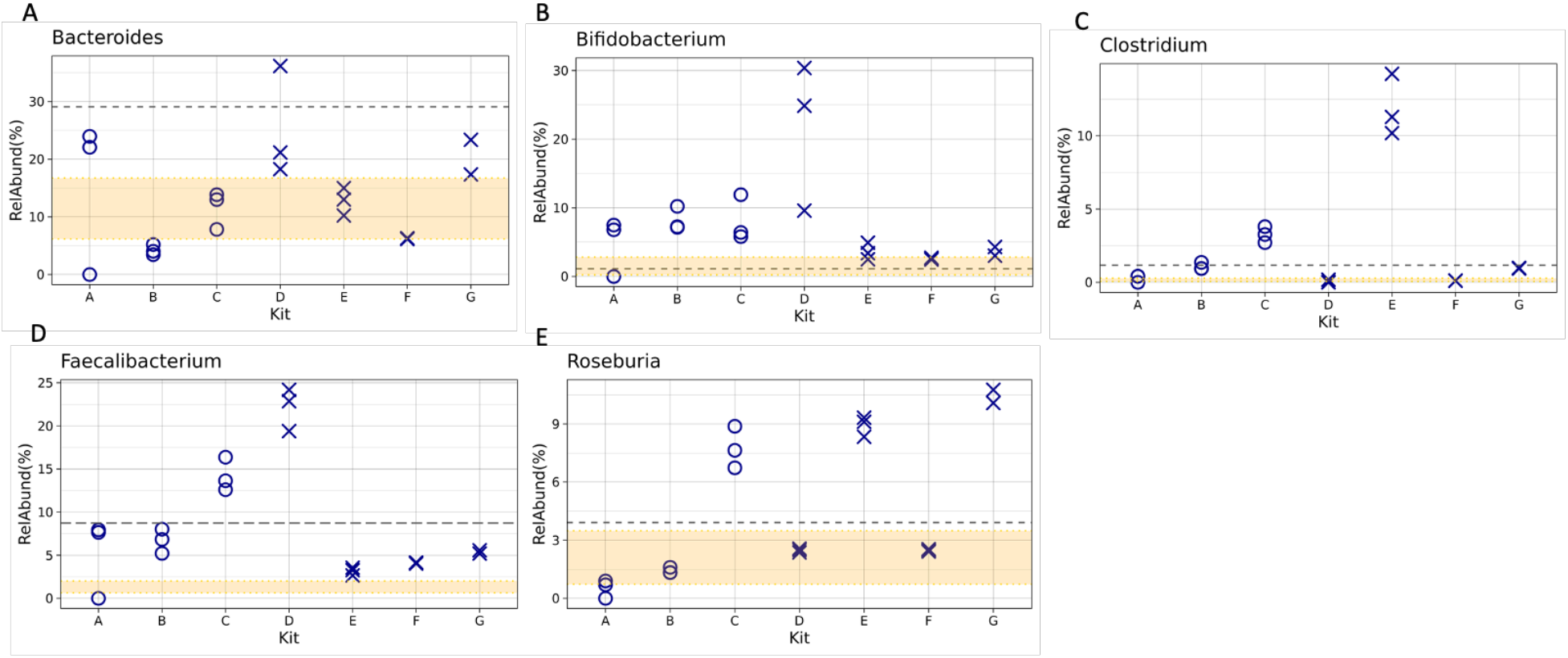
Comparison of Relative Abundance for Select Genera. Relative abundance for select genera **(A)** *Bacteroides* **(B)** *Bifidobacterium* **(C)** *Clostridium* (to include *Clostridioides*) **(D)** *Faecalibacterium* and **(E)** *Roseburia* were plotted compared to reported comparators from company G as a grey dashed line and from company E as a shaded yellow region. The grey dashed line represents the average from the Human Gut project reported by company G as a comparator. The yellow shaded region indicates the IQR used to define the healthy population as a comparator used by company E. The symbols indicated metagenomic analysis: O’s designate companies using 16S amplicon sequencing and X’s designate WMS analysis. Three replicates are shown for each Company except for Company F which only had 2 successful samples.

Replicate 1 from Company A had a significantly different taxonomic profile compared to the two other replicates from Company A and those from other companies (Figure 3B). This egregious anomaly prompted further investigation. Compared to replicates 1 and 2 from Company A, replicate 3 had approximately half the number of reads (186K compared to just over 400K). This might have contributed to its distinct profile, as only 6 genera were common between replicate 3 and replicates 1 and 2, corresponding to 10% of the total genera and abundance for these replicates. In contrast, replicates 1 and 2 shared 69 out of 70 and 71 genera, respectively. Notably, the other 28 genera in replicate 3 were not unique but were identified in at least one other replicate across all the companies. It is also worth noting that the percentage of unspecified reads in replicate 3 comprised over 50% of the sample, whereas the unspecified designation accounted for just over 20% in replicates 1 and 2. These discrepancies raise concerns not just because the profile was distinct from other replicates originating from the same sample, but also because it apparently passed all quality control checks from the company, resulting in a report being generated and sent back to the customer.

### Differences in Health Indicators Across Companies

In addition to providing customers with a taxonomic profile, many reports also include health metrics, such as comparing a customer’s specific taxonomic profile to that of a comparator group. Notably, the comparator group varies between companies. For instance, Company E uses an internally developed dataset from a “healthy” cohort, while other companies rely on external datasets, such as those from the Human Microbiome Project [16] or American Gut [9]. Beyond providing comparative data, some companies also identify beneficial or harmful microbes and suggest ways to modulate these microbes’ abundance to improve health. To examine how results vary between companies on a taxa-by-taxa basis, five clinically relevant genera were selected (Figure 3 **(A)** *Bacteroides* **(B)** *Bifidobacterium* **(C)** *Clostridium* (to include *Clostridioides*) **(D)** *Faecalibacterium* and **(E)** *Roseburia*). All results were compared to the criteria provided by two of the companies. One company reported average values of the 5 genera in their reference population (Figure 3, gray dotted line), and the other company reported an interquartile range (IQR) of their reference population (Figure 3, range between the 25% and 75% bounds shaded in yellow). Interestingly, except for *Bifidobacterium*, the average used by one company does not fall within the IQR used by the other company. Variability between companies is observed for all five genera, with no apparent correlation to library preparation (16S rRNA gene amplicon sequencing vs WMS). In addition, the precision from a single company varied based on taxa, with some taxa showing indistinguishable points between replicates (*e*.*g*., Figure 3E, *Roseburia* for companies D and F) and other showing wide variability (Figure 3B, *Bifidobacterium* for company D). The implications of this variability are particularly evident in the health recommendations commonly included in DTC microbiome testing reports. The most striking example is of the conflicting recommendations between replicate 3 from Company A and replicates 1 and 2. For instance, starch breakdown was assessed as a metric for gut health; replicate 3 scored below average, whereas replicates 1 and 2 scored above. Out of the 10 functional categories assessed by this company, seven were below average for replicate 3, compared to only one for replicates 1 and 2, and the below-average category for replicates 1 and 2 was distinct from the seven observed for replicate 3. The company reported the microbiome as healthy for replicate 1 and 2, but unhealthy for replicate 3, which could lead to unnecessary interventions if such a report was received by a consumer. While Company A provides an extreme example, other discrepancies, including the identification of pathogens, were observed between different companies. Notably, *Clostridioides difficile*, a primary causative agent of antibiotic-associated diarrhea for which carriage is a risk factor, was not consistently identified between companies.

## Discussion

Our rigorous assessment of seven microbiome testing companies has spotlighted the systemic issue of poor reproducibility that plagues the industry. The primary contributor to this outcome is methodological variability. The significant impact of methodological variables on the results of microbiome metagenomic measurements has been demonstrated by us and others, and DTC microbiome testing companies are not immune to these sources of bias [14, 17-20]. The use of a homogenous stool reference material enabled a systematic evaluation without the confounding factors of biological or composition heterogeneity that may be found in a traditional stool sample. Despite these benefits, it is important to emphasize the stool reference material used for this study does not represent an absolute “ground truth”. Consequently, this study cannot definitively determine which results are closest to the actual microbiome composition, nor was that the goal of this study. Nonetheless, access to a homogenous and standardized material, comparable to the complexity to a real-world sample, is sufficient for evaluating the precision of the sample processing workflows. Overall, this study revealed that the impact of methodological variability (between company comparisons) was on the same order of magnitude as biological variability (samples taken from 8 distinct donors). These findings have major implications for the comparability of tests on a single individual from two different companies and highlights the need for caution in interpreting and acting on these test results, especially in the absence of validated diagnostic evidence. Of note, despite the promising potential of the gut microbiome in human health, as evidenced by recent drug approvals [21-23], the diagnostic capabilities of these tests remain largely underdeveloped, partly due to the challenge demonstrated in this study.

Another crucial aspect we considered was the validity of the diagnostic and prognostic implications derived from these tests. Questions arise as to how a testing service determines whether a patient’s microbiome is healthy or unhealthy, what the relevant diagnostic and prognostic biomarkers are, and how these biomarkers were identified and validated. These questions are significant, as the test results may lead consumers to make potentially unwarranted or unsafe lifestyle changes. Many of the results are reported with respect to an average or “healthy” microbiome; however, defining a healthy microbiome remains a challenge due to the heterogeneity of the human population, confounders, and the possibility of multiple healthy microbiome definitions. Even among the DTC companies evaluated, there was no standard practice for selecting a comparative ‘heathy’ population. In some cases, companies used data generated in-house, either by measuring a cohort specifically recruited to serve as the comparative population or by aggregating the data from prior customer’s test samples. Alternatively, some companies look to previously constructed datasets such as that from American Gut or the Human Microbiome Project (HMP); both methods raise concerns. A benefit of using data generated in-house is that it was likely collected following the same workflow being used for the test samples, which should reduce the impact of methodological variability as reproducibility within a workflow is generally very good. However, little information was given on the demographics of these comparative populations, raising questions about how representative they are. In contrast, comparisons using data from previous efforts such as the American Gut, may provide more demographic information and/or generated a larger more representative dataset than possible for an individual company. As detailed above, methodological variability is a major hurdle for data comparability; therefore, if the DTC company is not following the same protocol used to generate the comparative population data, observed differences could be due either to biology, methodology, or a combination.

In general, the health recommendations of many DTC microbiome testing companies focus on healthier eating habits, which may seem innocuous and unlikely to cause direct harm to the consumers. However, it is crucial to recognize that a significant portion of the individuals who seek these tests are often those with chronic gut conditions, who may have struggled to find effective treatments. For these individuals, the variability in test results or the lack of expected outcomes from the recommendations provided, could not only lead to a loss of consumer faith in the science but also result in the delay of appropriate medical care. Additionally, some companies recommended that customers start taking costly supplements (e.g., probiotics) that are sold by the same company and for which there is very little clinical evidence for efficacy.

Given that adopting a single methodological workflow across the entire microbiome industry is unrealistic, there are several steps that could be taken to improve result transparency and interpretation. There are two primary areas to address with respect to DTC testing (1) clinical validity and (2) analytical performance. With respect to the first, the rapidly expanding field of microbiome research is constantly improving our understanding of the associations between health and the microbiome, but much of the data is likely correlative rather than causative. Without substantial causative evidence, the clinical validity of these tests will remain a challenge. While not a solution, inclusion of peer reviewed literature citations to support any claims to health status, or designation of a microbe as beneficial or harmful, would provide consumers (and clinicians) with a means of understanding the legitimacy of their results.

There are many more readily available resources for addressing analytical performance. An ideal standard which would enable both accurate and precise testing does not yet exist; however, as demonstrated by this study, inclusion of standard fecal material could be used to determine the minimum number of reads required for reproducible results and other minimum standards to ensure result validity. Additionally, while on a smaller scale than the 100s of organisms present in a stool sample, mock communities of 10-20 common gut microbes are available from several commercial outlets that enable end-users to assess the accuracy and precision of their microbiome measurements. Although beyond the scope of this discussion, in addition to the reference materials mentioned here, there are numerous publications on validating analytical performance of a metagenomic workflow [24-26]. The adoption of standards, and the reporting of the analytical performance gleaned from these standards, would help customers and other stakeholders (including regulatory bodies) understand and characterize bias in a given workflow. The industry should consider developing a consensus document on minimum requirements, with input from testing companies and other stakeholders. Similar documents, such as MIQE, EMMI, AsqMI, and STORMS have been developed for other high complexity molecular diagnostic assays [27-30]. Such a document would support test validity and bolster consumer confidence for DTC gut microbiome testing.

## Methods

### Material preparation

As part of the Microbiome Program, researchers at NIST have undertaken the challenge of producing a whole stool gut microbiome reference material. While the final reference material will be comprised of pooled donor material, as part of early investigations NIST obtained homogenized material from individual donors. Donor material was obtained from multiple volunteer donors by The BioCollective (Denver, CO, USA). All whole stool samples were collected after informed consent under approved IRB protocols at The BioCollective. All work conducted at NIST was reviewed and approved by the U. S. National Institute of Standards and Technology Research Protections Office. Having a relatively large quantity of homogenized stool gave NIST the confidence that any deviations seen in the results were due to bias introduced during the sample processing workflow and not biological differences between samples. To further ensure a homogenous starting sample, multiple aliquots were pooled in a single 50 mL conical tube, to a final volume of approximately 540 mg/mL stool and a Bristol stool consistency of 4, for sampling. This pooled material was used to inoculate all tests within 1 hour of preparation.

### NIST Analysis of the homogenized whole stool sample

In addition to the analyses provided by the service providers, the standardized fecal sample was also analyzed in the NIST lab using in-house DNA extraction, 16S rRNA and WMS workflows, to provide a better understanding of how reproducibility within a service provider compared to reproducibility within our own lab. Four replicate DNA extractions were carried out using the Zymo Research Quick-DNA Fecal/Soil microbe miniprep extraction kit [cat # D6010] per protocol with the vortex genie adapter for bead beating carried out for 45 mins. WMS and 16S rRNA sequencing library preps were carried out on the same DNA extract. 16S rRNA library prep was carried out as followed: 12 ng of genomic DNA was amplified in a 25 µL reaction comprised of 2x Kapa HiFi Master Mix (Roche cat # KK2601/2), 1 µL of 10 µM V4 515F/806R primers with Illumina adapters (5’-TCGTCGGCAGCGTCAGATGTGTATAAGAGACAGGTGYCAGCMGCCGCGGTAA -3’, 5’-GTCTCGTGGGCTCGGAGATGTGTATAAGAGACAGGGACTACNVGGGTWTCTAAT -3’), under the following conditions: 95.0 °C (3 min), and 18 cycles of amplification 98.0 °C (30 sec), 55 °C (15 sec), 72 °C (20 sec), and a final extraction at 72 °C for 5 min. 16S amplification products were purified with SPRIselect beads (Beckman Coulter cat # B23317/ B23318/ B23319) in a 0.8:1 (bead to product) ratio. The cleaned 16S products were indexed using IDT® for Illumina® DNA/RNA UD Indexes with 8 amplification cycles, but the same thermocycling conditions used for the 16S amplification. The indexed 16S amplicons were purified with SPRIselect beads in a 1:1 (bead to product) ratio. Purified indexed amplicons were pooled in equal quantities and adjusted to a 4nM pool for sequencing preparation following the MiSeq Systems Denature and Dilute DNA Libraries Guide, Protocol A (Document # 15039740 v10, February 2019).

WMS library prep was carried out using Covaris focused-ultrasonication for fragmentation followed by NEB for indexing. 10 µL of the DNA extract (approximately 500-700 ng of DNA) was added to a 120 µL of TE in a microtube AFA Fiber Pre-slit Snap Cap 6X16mm tube (Covaris Cat#: 520045). Samples were fragmented using the following settings: Intensity: 10%; Duty cycle: 10%; Cycle/burst (CPB): 100; Time: 1 min. Fragmented DNA was analyzed using the Tapestation and average peak size was approximately 280 bp. Fragmented DNA was indexed using the New England Biolab (NEB) NEBNext Ultra II DNA Library Prep Kit for Illumina with NEBNext Multiplex oligos for Illumina (NEB cat # E7645S/L; E6440) following the instruction manual (V6.1_5/20) with 50 µL of fragmented DNA as input; this corresponded to approximately 200 – 300 ng of DNA input. Purified indexed samples were pooled in equal quantities and adjusted to a 4nM pool for sequencing preparation following the MiSeq Systems Denature and Dilute DNA Libraries Guide, Protocol A (Document # 15039740 v10, February 2019). Both 16S and WMS sequencing libraries were run using a MiSeq Reagent Kit v3 600-cycle cartridge (MS-102-3003) with pair-end reads (raw reads will be available at https://doi.org/10.18434/mds2-3348; contact the link is not active).

16S reads were analyzed in R (4.1.0) using CutAdapt [31] for primer trimming and DADA2 (1.20.0) [32] with the silva database [33] (v138) for taxonomic classification. The complete R code is included in Supplemental file 1 as an R Markdown and html (16S_Analysis/Dada2-16sAnalysis2024).

WMS reads were taxonomically profiled using Kraken2 [34] and bracken [35] with default parameters against the prebuilt GTDB-tk database(r89) [36, 37]. Finally, read quality was assessed with FastQC. The integrated snakemake workflow, based on taxonomic profiling from NBIS-Meta (https://github.com/NBISweden/nbis-meta) can be found at https://github.com/ikeenum/WW_influent_bias.

### Analysis of additional donor samples

WMS was also conducted on the 7 additional donors of the candidate fecal reference material to serve as comparators for assessing the magnitude of methodological variability compared to biological (donor) variability. Two replicate DNA extractions from homogenized whole stool from all 8 unique donors (the donor used for all previous analyses plus 7 additional donors) was carried out using the Qiagen PowerSoil Pro Kit. WMS library preparation was done using Illumina NexteraXT followed by sequencing on the Illumina HiSeq X platform (raw reads will be available at https://doi.org/10.18434/mds2-3348; contact if the link is not active). WMS reads were profiled using the Kraken and bracken workflows as described above.

### Selection and preparation of DTC Gut Microbiome Tests

NIST conducted a web search for “at home direct to consumer microbiome testing kits” and selected 7 companies for inclusion in the study; 3 kits were ordered from each company to assess the intra-and inter-laboratory precision (reproducibility). A NIST employee created an account for each company using a personal email account so that the companies were blinded to the experiment at the time of testing.

Each test kit came with instructions, collection devices, sample containers, and pre-paid shippers. To the extent possible, the sampling instructions were followed per the manufacturers’ instructions. There were two notable deviations from the sampling instructions. First, some companies provided proprietary collection devices to support sampling directly from a bowel movement, while others instructed users to sample from used toilet paper; for this experiment all samples were taken directly from the 50mL conical. Second, some companies provided instructions to sample multiple sections on the bowel movement, which was not possible with our sample type. As mentioned above, all kits were inoculated within one hour of preparing the sample and then immediately packaged in the prepaid shippers and dropped off for shipping at the respective location (e.g. USPS, FedEx) as designated by the shipping label.

### Analysis of Taxonomic Profiles

Six of the 7 service providers produced a report of analysis for each kit (n=3) sent. One service provider produced 2 reports and reported a sample failure. An additional sample was sent, and this also failed their sample QC (the specific type of failure was not disclosed) and a third attempt was not made. Provider reports varied in content and format necessitating manual curation of the taxonomic data from each report to produce a consistent format. In addition to creating a consistent file and format, some datasets provided taxonomic resolution down to the species or even strain level whereas other taxa were identified at a higher order, for example only down to the family level. A genus level taxonomic classification was chosen to facilitate comparisons between datasets; therefore, the relative abundance of all species and strain level designations were summed to the genus level. The relative abundances for unclassified taxa and taxa where a specific genus designation was not given were summed and given an unspecified designation. Taxonomic tables generated from the NIST 16S and WMS analysis were put in the same format. All resulting tables were used as the input files for future analysis conducted in R Studio (Supplemental files 1 and 2, NIST_data and Genus_tables). We are unable to share the actual reports in a blinded fashion so this version of the data will not be made publicly available. The complete code for analysis and figure generation can be found in Supplemental file 3 as an R Markdown and html (2024-analysis).

## Disclaimer

Certain equipment, instruments, software, or materials are identified in this paper in order to specify the experimental procedure adequately. Such identification is not intended to imply recommendation or endorsement of any product or service by NIST, nor is it intended to imply that the materials or equipment identified are necessarily the best available for the purpose.

