## Supplemental File 1 for "Evaluating the Analytical Performance of Direct-to-Consumer Gut Microbiome Testing Services": Dada2-16sAnalysis2024.html

16S analysis


### 16S analysis

```
rm(list=ls())    # Clear variables; clear plots
```

('tidyverse','readxl','dada2','broom','rstudioapi','stringr', 'stats' , 'compiler', 'backports', 'magrittr', 'rprojroot', 'tools', 'htmltools', 'yaml', 'Rcpp', 'stringi', 'rmarkdown', 'knitr', 'digest', 'evaluate', 'seqinr', 'vegan')

```
sapply(c('tidyverse','readxl','dada2', 'rmarkdown', 'knitr', 'Rcpp'), require, character.only = TRUE) # Load packages
```

```
## Loading required package: tidyverse
```

```
## ── Attaching packages ─────────────────────────────────────── tidyverse 1.3.1 ──
```

```
## ✔ ggplot2 3.4.3     ✔ purrr   0.3.4
## ✔ tibble  3.1.2     ✔ dplyr   1.0.7
## ✔ tidyr   1.1.3     ✔ stringr 1.4.0
## ✔ readr   1.4.0     ✔ forcats 0.5.1
```

```
## ── Conflicts ────────────────────────────────────────── tidyverse_conflicts() ──
## ✖ dplyr::filter() masks stats::filter()
## ✖ dplyr::lag()    masks stats::lag()
```

```
## Loading required package: readxl
```

```
## Loading required package: dada2
```

```
## Loading required package: Rcpp
```

```
## Loading required package: rmarkdown
```

```
## Loading required package: knitr
```

```
## tidyverse    readxl     dada2 rmarkdown     knitr      Rcpp 
##      TRUE      TRUE      TRUE      TRUE      TRUE      TRUE
```

```
theme_set(theme_bw() + theme(plot.title = element_text(size = 11, face = 'bold', hjust = 0.5)))
sessionInfo()
```

```
## R version 4.1.0 (2021-05-18)
## Platform: x86_64-pc-linux-gnu (64-bit)
## Running under: Ubuntu 20.04.2 LTS
## 
## Matrix products: default
## BLAS:   /usr/lib/x86_64-linux-gnu/blas/libblas.so.3.9.0
## LAPACK: /usr/lib/x86_64-linux-gnu/lapack/liblapack.so.3.9.0
## 
## locale:
##  [1] LC_CTYPE=C.UTF-8       LC_NUMERIC=C           LC_TIME=C.UTF-8       
##  [4] LC_COLLATE=C.UTF-8     LC_MONETARY=C.UTF-8    LC_MESSAGES=C.UTF-8   
##  [7] LC_PAPER=C.UTF-8       LC_NAME=C              LC_ADDRESS=C          
## [10] LC_TELEPHONE=C         LC_MEASUREMENT=C.UTF-8 LC_IDENTIFICATION=C   
## 
## attached base packages:
## [1] stats     graphics  grDevices utils     datasets  methods   base     
## 
## other attached packages:
##  [1] knitr_1.33      rmarkdown_2.9   dada2_1.20.0    Rcpp_1.0.6     
##  [5] readxl_1.3.1    forcats_0.5.1   stringr_1.4.0   dplyr_1.0.7    
##  [9] purrr_0.3.4     readr_1.4.0     tidyr_1.1.3     tibble_3.1.2   
## [13] ggplot2_3.4.3   tidyverse_1.3.1
## 
## loaded via a namespace (and not attached):
##  [1] bitops_1.0-7                matrixStats_0.59.0         
##  [3] fs_1.5.0                    lubridate_1.7.10           
##  [5] RColorBrewer_1.1-2          httr_1.4.2                 
##  [7] GenomeInfoDb_1.28.0         tools_4.1.0                
##  [9] backports_1.2.1             utf8_1.2.1                 
## [11] R6_2.5.0                    DBI_1.1.1                  
## [13] BiocGenerics_0.38.0         colorspace_2.0-2           
## [15] withr_2.5.0                 tidyselect_1.1.1           
## [17] compiler_4.1.0              cli_3.6.1                  
## [19] rvest_1.0.0                 Biobase_2.52.0             
## [21] xml2_1.3.2                  DelayedArray_0.18.0        
## [23] scales_1.2.1                digest_0.6.27              
## [25] Rsamtools_2.8.0             XVector_0.32.0             
## [27] jpeg_0.1-8.1                pkgconfig_2.0.3            
## [29] htmltools_0.5.1.1           MatrixGenerics_1.4.0       
## [31] dbplyr_2.1.1                rlang_1.1.1                
## [33] rstudioapi_0.13             generics_0.1.0             
## [35] hwriter_1.3.2               jsonlite_1.7.2             
## [37] BiocParallel_1.26.0         RCurl_1.98-1.3             
## [39] magrittr_2.0.1              GenomeInfoDbData_1.2.6     
## [41] Matrix_1.3-3                munsell_0.5.0              
## [43] S4Vectors_0.30.0            fansi_0.5.0                
## [45] lifecycle_1.0.3             stringi_1.6.2              
## [47] yaml_2.2.1                  SummarizedExperiment_1.22.0
## [49] zlibbioc_1.38.0             plyr_1.8.6                 
## [51] grid_4.1.0                  parallel_4.1.0             
## [53] crayon_1.4.1                lattice_0.20-44            
## [55] Biostrings_2.60.1           haven_2.4.1                
## [57] hms_1.1.0                   pillar_1.6.1               
## [59] GenomicRanges_1.44.0        reshape2_1.4.4             
## [61] stats4_4.1.0                reprex_2.0.0               
## [63] glue_1.4.2                  evaluate_0.14              
## [65] ShortRead_1.50.0            latticeExtra_0.6-29        
## [67] RcppParallel_5.1.4          modelr_0.1.8               
## [69] vctrs_0.6.3                 png_0.1-7                  
## [71] cellranger_1.1.0            gtable_0.3.0               
## [73] assertthat_0.2.1            xfun_0.24                  
## [75] broom_0.7.8                 GenomicAlignments_1.28.0   
## [77] IRanges_2.26.0              ellipsis_0.3.2
```

```
print(getwd())
```

```
## [1] "/home/data/DTC-microbiome/publication_set"
```

NIST samples run on Illumina 2x300 w/V4 16s primers (515F/806R)

```
# V4 reads -- CUTADAPT

#FWD: GTGYCAGCMGCCGCGGTAA
#REV: GGACTACNVGGGTWTCTAAT

source("NIST_data/16S_Analysis/cutadapt_v4.R") 

forward.reads <- sort(list.files("NIST_data/16S_Analysis/raw_reads", pattern = "R1_001.fastq.gz", full.names = TRUE)) 

reverse.reads <- sort(list.files("NIST_data/16S_Analysis/raw_reads", pattern = "R2_001.fastq.gz", full.names = TRUE)) 

sample.names <- sapply(strsplit(basename(forward.reads), "_"), `[`, 1)

cutadapt("/usr/bin/cutadapt",
         output.directory = "NIST_data/16S_Analysis/primertrim_reads/",
         forward.primer = "^GTGYCAGCMGCCGCGGTAA",
         reverse.primer = "^GACTACNVGGGTWTCTAAT",
         error.rate = 0.15,
         minumum.overlap = 19,
         clip.from.start.read1 = 0,
         clip.from.start.read2 = 1,
         maximum.N = 0,
         discard.untrimmed = TRUE,
         filter.pairs = "any",
         log.files = TRUE,
         R.cores = 4 )

save.image("NIST-16SAnalysis.RData")
```

```
plotQualityProfile(forward.reads[1:4])
```

```
## Warning: The `<scale>` argument of `guides()` cannot be `FALSE`. Use "none"
## instead as of ggplot2 3.3.4.
```

```
plotQualityProfile(reverse.reads[1:4])
```

```
filtFs <- file.path("NIST_data/16S_Analysis/Qtrim_reads", paste0(sample.names, "_R1_filtered.fastq.gz"))

filtRs <- file.path("NIST_data/16S_Analysis/Qtrim_reads", paste0(sample.names, "_R2_filtered.fastq.gz"))

out <- filterAndTrim(forward.reads, filtFs, reverse.reads, filtRs, 
                     truncLen = c(275, 250),
                     maxN = c(0, 0), 
                     maxEE = c(5, 5),
                     rm.phix = TRUE,
                     compress = TRUE, 
                     multithread = TRUE)

out %>% as.data.frame() %>%
  dplyr::mutate(percent.surviving = round(reads.out/reads.in*100, 1))
```

```
##                                reads.in reads.out percent.surviving
## NIST-1_S5_L001_R1_001.fastq.gz   679069    652828              96.1
## NIST-2_S6_L001_R1_001.fastq.gz   552272    532874              96.5
## NIST-3_S7_L001_R1_001.fastq.gz   614937    593559              96.5
## NIST-4_S8_L001_R1_001.fastq.gz   865043    839490              97.0
```

```
save.image("NIST-16SAnalysis.RData")
```

```
errF <- learnErrors(filtFs, multithread = TRUE, randomize = TRUE, nbases = 2e8, verbose = TRUE)
```

```
## 377400100 total bases in 1372364 reads from 2 samples will be used for learning the error rates.
## Initializing error rates to maximum possible estimate.
## selfConsist step 1 ..
##    selfConsist step 2
##    selfConsist step 3
##    selfConsist step 4
##    selfConsist step 5
##    selfConsist step 6
##    selfConsist step 7
## Convergence after  7  rounds.
```

```
errR <- learnErrors(filtRs, multithread = TRUE, randomize = TRUE, nbases = 2e8, verbose = TRUE)
```

```
## 358262250 total bases in 1433049 reads from 2 samples will be used for learning the error rates.
## Initializing error rates to maximum possible estimate.
## selfConsist step 1 ..
##    selfConsist step 2
##    selfConsist step 3
##    selfConsist step 4
##    selfConsist step 5
##    selfConsist step 6
##    selfConsist step 7
##    selfConsist step 8
##    selfConsist step 9
##    selfConsist step 10
```

```
## Self-consistency loop terminated before convergence.
```

```
save.image("NIST-16SAnalysis.RData")
```

```
plotErrors(errR, nominalQ = TRUE)
```

```
## Warning: Transformation introduced infinite values in continuous y-axis
```

```
plotErrors(errF, nominalQ = TRUE)
```

```
## Warning: Transformation introduced infinite values in continuous y-axis
```

```
derepFs <- derepFastq(filtFs, verbose = T)
```

```
## Dereplicating sequence entries in Fastq file: NIST_data/16S_Analysis/Qtrim_reads/NIST-1_R1_filtered.fastq.gz
```

```
## Encountered 66636 unique sequences from 652828 total sequences read.
```

```
## Dereplicating sequence entries in Fastq file: NIST_data/16S_Analysis/Qtrim_reads/NIST-2_R1_filtered.fastq.gz
```

```
## Encountered 56026 unique sequences from 532874 total sequences read.
```

```
## Dereplicating sequence entries in Fastq file: NIST_data/16S_Analysis/Qtrim_reads/NIST-3_R1_filtered.fastq.gz
```

```
## Encountered 60901 unique sequences from 593559 total sequences read.
```

```
## Dereplicating sequence entries in Fastq file: NIST_data/16S_Analysis/Qtrim_reads/NIST-4_R1_filtered.fastq.gz
```

```
## Encountered 82552 unique sequences from 839490 total sequences read.
```

```
names(derepFs) <- sample.names

save.image("NIST-16SAnalysis.RData")

derepRs <- derepFastq(filtRs, verbose = T)
```

```
## Dereplicating sequence entries in Fastq file: NIST_data/16S_Analysis/Qtrim_reads/NIST-1_R2_filtered.fastq.gz
```

```
## Encountered 145067 unique sequences from 652828 total sequences read.
```

```
## Dereplicating sequence entries in Fastq file: NIST_data/16S_Analysis/Qtrim_reads/NIST-2_R2_filtered.fastq.gz
```

```
## Encountered 118438 unique sequences from 532874 total sequences read.
```

```
## Dereplicating sequence entries in Fastq file: NIST_data/16S_Analysis/Qtrim_reads/NIST-3_R2_filtered.fastq.gz
```

```
## Encountered 144368 unique sequences from 593559 total sequences read.
```

```
## Dereplicating sequence entries in Fastq file: NIST_data/16S_Analysis/Qtrim_reads/NIST-4_R2_filtered.fastq.gz
```

```
## Encountered 181872 unique sequences from 839490 total sequences read.
```

```
names(derepRs) <- sample.names

save.image("NIST-16SAnalysis.RData")
```

```
dadaFs <- dada(derepFs, err = errF, multithread = TRUE)
```

```
## Sample 1 - 652828 reads in 66636 unique sequences.
## Sample 2 - 532874 reads in 56026 unique sequences.
## Sample 3 - 593559 reads in 60901 unique sequences.
## Sample 4 - 839490 reads in 82552 unique sequences.
```

```
save.image("NIST-16SAnalysis.RData")

dadaRs <- dada(derepRs, err = errR, multithread = TRUE)
```

```
## Sample 1 - 652828 reads in 145067 unique sequences.
## Sample 2 - 532874 reads in 118438 unique sequences.
## Sample 3 - 593559 reads in 144368 unique sequences.
## Sample 4 - 839490 reads in 181872 unique sequences.
```

```
save.image("NIST-16SAnalysis.RData")
```

```
mergers <- mergePairs(dadaFs, derepFs, dadaRs, derepRs, verbose = TRUE)
```

```
## 640390 paired-reads (in 13506 unique pairings) successfully merged out of 647507 (in 15756 pairings) input.
```

```
## 521892 paired-reads (in 12423 unique pairings) successfully merged out of 527965 (in 14324 pairings) input.
```

```
## 581517 paired-reads (in 12813 unique pairings) successfully merged out of 588657 (in 15062 pairings) input.
```

```
## 824747 paired-reads (in 15389 unique pairings) successfully merged out of 833369 (in 18230 pairings) input.
```

```
save.image("NIST-16SAnalysis.RData")
```

```
seqtab<- makeSequenceTable(mergers)
```

```
getN <- function(x) sum(getUniques(x))
NIST_track <- cbind(out, sapply(dadaFs, getN), sapply(dadaRs, getN), sapply(mergers, getN), rowSums(seqtab))
colnames(NIST_track) <- c("input", "filtered", "denoisedF", "denoisedR", "merged", "nonchimeric")
rownames(NIST_track) <- sample.names
NIST_track
```

```
##         input filtered denoisedF denoisedR merged nonchimeric
## NIST-1 679069   652828    651298    648656 640390      640390
## NIST-2 552272   532874    531262    529254 521892      521892
## NIST-3 614937   593559    592180    589770 581517      581517
## NIST-4 865043   839490    837726    834765 824747      824747
```

```
write.table(NIST_track, "NIST_data/16S_Analysis/dada2_output/NIST_Track.txt",
            quote = FALSE, row.names = TRUE, sep = "\t")

save.image("NIST-16SAnalysis.RData")
```

```
seqtab.nochim <- removeBimeraDenovo(seqtab, method="consensus", multithread=TRUE, verbose = TRUE)
```

```
## Identified 15707 bimeras out of 17755 input sequences.
```

```
asvtab <- t(as.data.frame(seqtab.nochim)) %>%
  as.data.frame() %>%
  tibble::rownames_to_column() %>%
  dplyr::mutate(asvID = paste0("asv", 1:nrow(.))) %>%
  plyr::rename(c("rowname" = "Sequence"))

save.image("NIST-16SAnalysis.RData")
```

```
taxa <- assignTaxonomy(seqtab.nochim, "NIST_data/16S_Analysis/silva_nr99_v138_wSpecies_train_set.fa.gz",
                       multithread=TRUE, tryRC = TRUE) %>%
  as.data.frame() %>%
  mutate(Sequence = rownames(.)) 
  
Taxa.tab <- full_join(taxa, asvtab) %>%
  gather(value = Count, key = Sample, -asvID, -Kingdom, -Phylum, -Class, -Order, -Family, -Genus, -Species, -Sequence) %>%
  group_by(Sample) %>%
  mutate(RelAbund = Count/sum(Count))
```

```
## Joining, by = "Sequence"
```

```
write.table(Taxa.tab, "NIST_data/16S_Analysis/dada2_output/taxa_tab.txt", quote = FALSE, row.names = FALSE, sep = "\t")

Taxa.tab2 <- Taxa.tab %>%
  select(-Count) %>%
  spread(value = "RelAbund", key = "Sample")

write.table(Taxa.tab2, "NIST_data/16S_Analysis/dada2_output/taxa_tab2.txt",
            quote = FALSE, row.names = FALSE, sep = "\t")
```
