## Supplemental File 1 for "Evaluating the Analytical Performance of Direct-to-Consumer Gut Microbiome Testing Services": Dada2-16sAnalysis2024.Rmd

````
---
title: "16S analysis"
output: html_document
---

```{r clear_workspace}
rm(list=ls())    # Clear variables; clear plots
```

('tidyverse','readxl','dada2','broom','rstudioapi','stringr', 'stats' , 'compiler', 'backports', 'magrittr', 'rprojroot', 'tools', 'htmltools', 'yaml', 'Rcpp', 'stringi', 'rmarkdown', 'knitr', 'digest', 'evaluate',  'seqinr', 'vegan')

```{r load_packages}
sapply(c('tidyverse','readxl','dada2', 'rmarkdown', 'knitr', 'Rcpp'), require, character.only = TRUE) # Load packages
theme_set(theme_bw() + theme(plot.title = element_text(size = 11, face = 'bold', hjust = 0.5)))
sessionInfo() 
print(getwd())
```
NIST samples run on Illumina 2x300 w/V4 16s primers (515F/806R)
```{r CUTADAPT_16S_primer_trimming}

# V4 reads -- CUTADAPT

#FWD: GTGYCAGCMGCCGCGGTAA
#REV: GGACTACNVGGGTWTCTAAT

source("NIST_data/16S_Analysis/cutadapt_v4.R") 

forward.reads <- sort(list.files("NIST_data/16S_Analysis/raw_reads", pattern = "R1_001.fastq.gz", full.names = TRUE)) 

reverse.reads <- sort(list.files("NIST_data/16S_Analysis/raw_reads", pattern = "R2_001.fastq.gz", full.names = TRUE)) 

sample.names <- sapply(strsplit(basename(forward.reads), "_"), `[`, 1)

cutadapt("/usr/bin/cutadapt",
         output.directory = "NIST_data/16S_Analysis/primertrim_reads/",
         forward.primer = "^GTGYCAGCMGCCGCGGTAA",
         reverse.primer = "^GACTACNVGGGTWTCTAAT",
         error.rate = 0.15,
         minumum.overlap = 19,
         clip.from.start.read1 = 0,
         clip.from.start.read2 = 1,
         maximum.N = 0,
         discard.untrimmed = TRUE,
         filter.pairs = "any",
         log.files = TRUE,
         R.cores = 4 )

save.image("NIST-16SAnalysis.RData")
```

```{r plot_quality_profile}
plotQualityProfile(forward.reads[1:4])          
plotQualityProfile(reverse.reads[1:4])
```

```{r DADA2_fliterAndTrim}

filtFs <- file.path("NIST_data/16S_Analysis/Qtrim_reads", paste0(sample.names, "_R1_filtered.fastq.gz"))

filtRs <- file.path("NIST_data/16S_Analysis/Qtrim_reads", paste0(sample.names, "_R2_filtered.fastq.gz"))

out <- filterAndTrim(forward.reads, filtFs, reverse.reads, filtRs, 
                     truncLen = c(275, 250),
                     maxN = c(0, 0), 
                     maxEE = c(5, 5),
                     rm.phix = TRUE,
                     compress = TRUE, 
                     multithread = TRUE)

out %>% as.data.frame() %>%
  dplyr::mutate(percent.surviving = round(reads.out/reads.in*100, 1))

save.image("NIST-16SAnalysis.RData")
```

```{r DADA2_learnError}
errF <- learnErrors(filtFs, multithread = TRUE, randomize = TRUE, nbases = 2e8, verbose = TRUE)
errR <- learnErrors(filtRs, multithread = TRUE, randomize = TRUE, nbases = 2e8, verbose = TRUE)
save.image("NIST-16SAnalysis.RData")
```

```{r DADA2_plotErrors}
plotErrors(errR, nominalQ = TRUE)
plotErrors(errF, nominalQ = TRUE)
```

```{r DADA2_derepFastq}
derepFs <- derepFastq(filtFs, verbose = T)
names(derepFs) <- sample.names

save.image("NIST-16SAnalysis.RData")

derepRs <- derepFastq(filtRs, verbose = T)
names(derepRs) <- sample.names

save.image("NIST-16SAnalysis.RData")
```

```{r DADA2_dada_sequence_error_correction}
dadaFs <- dada(derepFs, err = errF, multithread = TRUE)
save.image("NIST-16SAnalysis.RData")

dadaRs <- dada(derepRs, err = errR, multithread = TRUE)
save.image("NIST-16SAnalysis.RData")
```

```{r DADA2_mergePairs}
mergers <- mergePairs(dadaFs, derepFs, dadaRs, derepRs, verbose = TRUE) 
save.image("NIST-16SAnalysis.RData")
```

```{r DADA2_makeSequenceTable}
seqtab<- makeSequenceTable(mergers)
```

```{r make_table_tracking_reads}
getN <- function(x) sum(getUniques(x))
NIST_track <- cbind(out, sapply(dadaFs, getN), sapply(dadaRs, getN), sapply(mergers, getN), rowSums(seqtab))
colnames(NIST_track) <- c("input", "filtered", "denoisedF", "denoisedR", "merged", "nonchimeric")
rownames(NIST_track) <- sample.names
NIST_track

write.table(NIST_track, "NIST_data/16S_Analysis/dada2_output/NIST_Track.txt",
            quote = FALSE, row.names = TRUE, sep = "\t")

save.image("NIST-16SAnalysis.RData")
```

```{r DADA2_readRDS_removeBimera_asvtab}
seqtab.nochim <- removeBimeraDenovo(seqtab, method="consensus", multithread=TRUE, verbose = TRUE)

asvtab <- t(as.data.frame(seqtab.nochim)) %>%
  as.data.frame() %>%
  tibble::rownames_to_column() %>%
  dplyr::mutate(asvID = paste0("asv", 1:nrow(.))) %>%
  plyr::rename(c("rowname" = "Sequence"))

save.image("NIST-16SAnalysis.RData")
```

```{r Taxonomy_assignment}
taxa <- assignTaxonomy(seqtab.nochim, "NIST_data/16S_Analysis/silva_nr99_v138_wSpecies_train_set.fa.gz",
                       multithread=TRUE, tryRC = TRUE) %>%
  as.data.frame() %>%
  mutate(Sequence = rownames(.)) 
  
Taxa.tab <- full_join(taxa, asvtab, by = "Sequence") %>%
  gather(value = Count, key = Sample, -asvID, -Kingdom, -Phylum, -Class, -Order, -Family, -Genus, -Species, -Sequence) %>%
  group_by(Sample) %>%
  mutate(RelAbund = Count/sum(Count)) 

write.table(Taxa.tab, "NIST_data/16S_Analysis/dada2_output/taxa_tab.txt", quote = FALSE, row.names = FALSE, sep = "\t")

Taxa.tab2 <- Taxa.tab %>%
  select(-Count) %>%
  spread(value = "RelAbund", key = "Sample")

write.table(Taxa.tab2, "NIST_data/16S_Analysis/dada2_output/taxa_tab2.txt",
            quote = FALSE, row.names = FALSE, sep = "\t")
```
````
