## Supplemental File 3 for "Evaluating the Analytical Performance of Direct-to-Consumer Gut Microbiome Testing Services": 2024-analysis.html

Comparison of Microbiome Profiles from DTC testing companies


### Comparison of Microbiome Profiles from DTC testing companies

```
rm(list=ls())    # Clear variables; clear plots
```

```
sapply(c('plyr', 'tidyverse','readxl',  'rmarkdown', 'knitr', 'vegan', 'ggforce','stringr'), require, character.only = TRUE) # Load packages
```

```
## Loading required package: plyr
```

```
## Loading required package: tidyverse
```

```
## ── Attaching packages ─────────────────────────────────────── tidyverse 1.3.1 ──
```

```
## ✔ ggplot2 3.4.3     ✔ purrr   0.3.4
## ✔ tibble  3.1.2     ✔ dplyr   1.0.7
## ✔ tidyr   1.1.3     ✔ stringr 1.4.0
## ✔ readr   1.4.0     ✔ forcats 0.5.1
```

```
## ── Conflicts ────────────────────────────────────────── tidyverse_conflicts() ──
## ✖ dplyr::arrange()   masks plyr::arrange()
## ✖ purrr::compact()   masks plyr::compact()
## ✖ dplyr::count()     masks plyr::count()
## ✖ dplyr::failwith()  masks plyr::failwith()
## ✖ dplyr::filter()    masks stats::filter()
## ✖ dplyr::id()        masks plyr::id()
## ✖ dplyr::lag()       masks stats::lag()
## ✖ dplyr::mutate()    masks plyr::mutate()
## ✖ dplyr::rename()    masks plyr::rename()
## ✖ dplyr::summarise() masks plyr::summarise()
## ✖ dplyr::summarize() masks plyr::summarize()
```

```
## Loading required package: readxl
```

```
## Loading required package: rmarkdown
```

```
## Loading required package: knitr
```

```
## Loading required package: vegan
```

```
## Loading required package: permute
```

```
## Loading required package: lattice
```

```
## This is vegan 2.6-4
```

```
## Loading required package: ggforce
```

```
##      plyr tidyverse    readxl rmarkdown     knitr     vegan   ggforce   stringr 
##      TRUE      TRUE      TRUE      TRUE      TRUE      TRUE      TRUE      TRUE
```

```
theme_set(theme_bw() + theme(plot.title = element_text(size = 11, face = 'bold', hjust = 0.5)))
sessionInfo()
```

```
## R version 4.1.0 (2021-05-18)
## Platform: x86_64-pc-linux-gnu (64-bit)
## Running under: Ubuntu 20.04.2 LTS
## 
## Matrix products: default
## BLAS:   /usr/lib/x86_64-linux-gnu/blas/libblas.so.3.9.0
## LAPACK: /usr/lib/x86_64-linux-gnu/lapack/liblapack.so.3.9.0
## 
## locale:
##  [1] LC_CTYPE=C.UTF-8       LC_NUMERIC=C           LC_TIME=C.UTF-8       
##  [4] LC_COLLATE=C.UTF-8     LC_MONETARY=C.UTF-8    LC_MESSAGES=C.UTF-8   
##  [7] LC_PAPER=C.UTF-8       LC_NAME=C              LC_ADDRESS=C          
## [10] LC_TELEPHONE=C         LC_MEASUREMENT=C.UTF-8 LC_IDENTIFICATION=C   
## 
## attached base packages:
## [1] stats     graphics  grDevices utils     datasets  methods   base     
## 
## other attached packages:
##  [1] ggforce_0.4.1   vegan_2.6-4     lattice_0.20-44 permute_0.9-7  
##  [5] knitr_1.33      rmarkdown_2.9   readxl_1.3.1    forcats_0.5.1  
##  [9] stringr_1.4.0   dplyr_1.0.7     purrr_0.3.4     readr_1.4.0    
## [13] tidyr_1.1.3     tibble_3.1.2    ggplot2_3.4.3   tidyverse_1.3.1
## [17] plyr_1.8.6     
## 
## loaded via a namespace (and not attached):
##  [1] Rcpp_1.0.6        lubridate_1.7.10  assertthat_0.2.1  digest_0.6.27    
##  [5] utf8_1.2.1        R6_2.5.0          cellranger_1.1.0  backports_1.2.1  
##  [9] reprex_2.0.0      evaluate_0.14     httr_1.4.2        pillar_1.6.1     
## [13] rlang_1.1.1       rstudioapi_0.13   Matrix_1.3-3      splines_4.1.0    
## [17] polyclip_1.10-4   munsell_0.5.0     broom_0.7.8       compiler_4.1.0   
## [21] modelr_0.1.8      xfun_0.24         pkgconfig_2.0.3   mgcv_1.8-35      
## [25] htmltools_0.5.1.1 tidyselect_1.1.1  fansi_0.5.0       crayon_1.4.1     
## [29] dbplyr_2.1.1      withr_2.5.0       MASS_7.3-54       grid_4.1.0       
## [33] nlme_3.1-152      jsonlite_1.7.2    gtable_0.3.0      lifecycle_1.0.3  
## [37] DBI_1.1.1         magrittr_2.0.1    scales_1.2.1      cli_3.6.1        
## [41] stringi_1.6.2     farver_2.1.0      fs_1.5.0          xml2_1.3.2       
## [45] ellipsis_0.3.2    generics_0.1.0    vctrs_0.6.3       tools_4.1.0      
## [49] glue_1.4.2        tweenr_2.0.2      hms_1.1.0         parallel_4.1.0   
## [53] yaml_2.2.1        colorspace_2.0-2  cluster_2.1.2     rvest_1.0.0      
## [57] haven_2.4.1
```

```
print(getwd())
```

```
## [1] "/home/data/DTC-microbiome/publication_set"
```

```
#to load workspace generated at NIST
#load("NIST-analysis.RData")
```

this analysis includes samples analyzed completely by NIST and the results from the DTC testing companies. The re-analysis of DTC by NIST is not included. Full data can be foudn in datasheet-build-070623

metadata

```
Meta <- read_excel("meta.xlsx")
```

```
pal_sample <- c("red", "gold", "green3", "purple2", "navy", "gray",
                "turquoise", "hotpink", "springgreen", "coral2", "purple", "powderblue",
                "orange", "lightslateblue", "lightseagreen", "deeppink", "orangered","black")

pal_35 <- c("#1F78C8" ,"#ff0000", "#33a02c" ,"#6A33C2" ,"#ff7f00", 
            "#565656","#FFD700","#a6cee3", "#FB6496" ,"#b2df8a", 
            "#CAB2D6","#FDBF6F","#0000FF","#EEE685", "#C8308C", 
            "#FF83FA" ,"#C814FA", "#999999" , "#36648B", "#00E2E5",
            "#00FF00", "#778B00", "#BEBE00","#8B3B00","#A52A3C" ,
            "#F0A0FF" ,"#0075DC", "#993F00" ,"#4C005C","#191919", 
            "#005C31", "#2BCE48" ,"#FFCC99" ,"#808080", "#94FFB5")

#color descriptors 
#1F78C8: A shade of blue, #ff0000: Red, #33a02c: Green, #6A33C2: Purple, #ff7f00: Orange, #565656: Dark gray, #FFD700: Gold/yellow, #a6cee3: Light blue, #FB6496: Pink, #b2df8a: Light green, #CAB2D6: Light purple, #FDBF6F: Light orange, #0000FF: Blue, #EEE685: Light yellow, #C8308C: Dark pink, #FF83FA: Light purple, #C814FA: Dark purple, #999999: Gray,, #36648B: Dark blue, #00E2E5: Light blue, #00FF00: Bright green, #778B00: Olive green, #BEBE00: Dark yellow, #8B3B00: Dark orange, #A52A3C: Dark red, #F0A0FF: Light pink, #0075DC: Bright blue, #993F00: Dark brown, #4C005C: Dark purple, #191919: Very dark gray/black, #005C31: Dark green, #2BCE48: Light green, #FFCC99: Light peach, #808080: Medium gray, #94FFB5: Light greenish-blue
```

### DTC data import

### DTC data - already formated as Genus Table needed for diversity metrics

```
Genus.A1 <- read_excel("Genus_tables/Genus_A.xlsx", 1) %>%
  group_by(Genus) %>%
  summarise_each(funs(sum))
```

```
## Warning: `summarise_each_()` was deprecated in dplyr 0.7.0.

## Warning: Please use `across()` instead.
```

```
## Warning: `funs()` was deprecated in dplyr 0.8.0.

## Warning: Please use a list of either functions or lambdas: 
## 
##   # Simple named list: 
##   list(mean = mean, median = median)
## 
##   # Auto named with `tibble::lst()`: 
##   tibble::lst(mean, median)
## 
##   # Using lambdas
##   list(~ mean(., trim = .2), ~ median(., na.rm = TRUE))
```

```
Genus.A2 <- read_excel("Genus_tables/Genus_A.xlsx", 2) %>%
  group_by(Genus) %>%
  summarise_each(funs(sum))
Genus.A3 <- read_excel("Genus_tables/Genus_A.xlsx", 3)%>%
  group_by(Genus) %>%
  summarise_each(funs(sum))
Genus_A <- full_join(Genus.A1, Genus.A2, by = "Genus") %>%
  full_join(., Genus.A3, by = "Genus") 
Genus_A %>%
  summarise_if(is.numeric, sum, na.rm = TRUE)
```

```
## # A tibble: 1 x 3
##   `A-1` `A-2` `A-3`
##   <dbl> <dbl> <dbl>
## 1     1  1.00     1
```

```
Genus.B1 <- read_excel("Genus_tables/Genus_B.xlsx", 1) %>%
  group_by(Genus) %>%
  summarise_each(funs(sum))
Genus.B2 <- read_excel("Genus_tables/Genus_B.xlsx", 2) %>%
  group_by(Genus) %>%
  summarise_each(funs(sum))
Genus.B3 <- read_excel("Genus_tables/Genus_B.xlsx", 3)%>%
  group_by(Genus) %>%
  summarise_each(funs(sum))
Genus_B <- full_join(Genus.B1, Genus.B2, by = "Genus") %>%
  full_join(., Genus.B3, by = "Genus")
Genus_B %>%
  summarise_if(is.numeric, sum, na.rm = TRUE)
```

```
## # A tibble: 1 x 3
##   `B-1` `B-2` `B-3`
##   <dbl> <dbl> <dbl>
## 1     1     1     1
```

```
Genus.C1 <- read_excel("Genus_tables/Genus_C.xlsx", 1)
Genus.C2 <- read_excel("Genus_tables/Genus_C.xlsx", 2)
Genus.C3 <- read_excel("Genus_tables/Genus_C.xlsx", 3)
Genus_C <- full_join(Genus.C1, Genus.C2, by = "Genus") %>%
  full_join(., Genus.C3, by = "Genus")
Genus_C %>%
  summarise_if(is.numeric, sum, na.rm = TRUE)
```

```
## # A tibble: 1 x 3
##   `C-1` `C-2` `C-3`
##   <dbl> <dbl> <dbl>
## 1     1     1     1
```

```
Genus.D1 <- read_excel("Genus_tables/Genus_D.xlsx", 1)%>%
  group_by(Genus) %>%
  summarise_each(funs(sum))
Genus.D2 <- read_excel("Genus_tables/Genus_D.xlsx", 2)%>%
  group_by(Genus) %>%
  summarise_each(funs(sum))
Genus.D3<- read_excel("Genus_tables/Genus_D.xlsx", 3)%>%
  group_by(Genus) %>%
  summarise_each(funs(sum))
Genus_D <- full_join(Genus.D1, Genus.D2, by = "Genus") %>%
  full_join(., Genus.D3, by = "Genus")
Genus_D %>%
  summarise_if(is.numeric, sum, na.rm = TRUE)
```

```
## # A tibble: 1 x 3
##   `D-1` `D-2` `D-3`
##   <dbl> <dbl> <dbl>
## 1  1.03 0.998 0.999
```

```
Genus.E1 <- read_excel("Genus_tables/Genus_E.xlsx", 1)
Genus.E2 <- read_excel("Genus_tables/Genus_E.xlsx", 2)
Genus.E3 <- read_excel("Genus_tables/Genus_E.xlsx", 3)
Genus_E <- full_join(Genus.E1, Genus.E2, by = "Genus") %>%
  full_join(., Genus.E3, by = "Genus")
Genus_E %>%
  summarise_if(is.numeric, sum, na.rm = TRUE)
```

```
## # A tibble: 1 x 3
##   `E-1` `E-2` `E-3`
##   <dbl> <dbl> <dbl>
## 1     1     1     1
```

```
Genus.F1 <- read_excel("Genus_tables/Genus_F.xlsx", 1)
Genus.F2 <- read_excel("Genus_tables/Genus_F.xlsx", 2)
Genus.F3 <- read_excel("Genus_tables/Genus_F.xlsx", 3)
Genus_F <- full_join(Genus.F1, Genus.F2, by = "Genus") %>%
  full_join(., Genus.F3, by = "Genus")
Genus_F %>%
  summarise_if(is.numeric, sum, na.rm = TRUE)
```

```
## # A tibble: 1 x 3
##   `F-1` `F-2` `F-3`
##   <dbl> <dbl> <dbl>
## 1     1  1.00     1
```

```
Genus.G1 <- read_excel("Genus_tables/Genus_G.xlsx", 1) %>%
  group_by(Genus) %>%
  summarise_each(funs(sum))
Genus.G2 <- read_excel("Genus_tables/Genus_G.xlsx", 2) %>%
  group_by(Genus) %>%
  summarise_each(funs(sum))
Genus_G <- full_join(Genus.G1, Genus.G2, by = "Genus") 
Genus_G[Genus_G == "Unclassified"] <- "Unspecified"
Genus_G %>%
  summarise_if(is.numeric, sum, na.rm = TRUE)
```

```
## # A tibble: 1 x 2
##   `G-1` `G-2`
##   <dbl> <dbl>
## 1  1.00  1.00
```

```
DTC.Genus.table <- full_join(Genus_A, Genus_B, by = "Genus") %>%
  full_join(., Genus_C, by = "Genus") %>%
  full_join(., Genus_D, by = "Genus") %>%
  full_join(., Genus_E, by = "Genus") %>%
  full_join(., Genus_F, by = "Genus") %>%
  full_join(., Genus_G, by = "Genus")

DTC.Genus.table %>%
  summarise_if(is.numeric, sum, na.rm = TRUE)
```

```
## # A tibble: 1 x 20
##   `A-1` `A-2` `A-3` `B-1` `B-2` `B-3` `C-1` `C-2` `C-3` `D-1` `D-2` `D-3` `E-1`
##   <dbl> <dbl> <dbl> <dbl> <dbl> <dbl> <dbl> <dbl> <dbl> <dbl> <dbl> <dbl> <dbl>
## 1     1  1.00     1     1     1     1     1     1     1  1.03 0.998 0.999     1
## # … with 7 more variables: E-2 <dbl>, E-3 <dbl>, F-1 <dbl>, F-2 <dbl>,
## #   F-3 <dbl>, G-1 <dbl>, G-2 <dbl>
```

```
#Long version of Genus.table
RA_DTCData <- DTC.Genus.table %>%
  gather(key = Sample, value = RelAbund, -Genus) %>%
  select(Sample, Genus, RelAbund) %>%
  mutate(Analysis = "DTC") %>%
  replace(is.na(.), 0) %>%
  mutate(Genus = str_replace(Genus,"unclassified", "Unspecified"))

#update location if running new analysis
save.image("NIST-analysis.RData")
```

### import NIST samples

```
#Shotgun data analyzed with Bracken
NIST5 <- read_excel("NIST_data/Bracken_reports.xlsx", 1) %>%
  mutate("Sample" = "NIST-5") %>%
  filter("new_est_reads" > 0)
NIST6  <- read_excel("NIST_data/Bracken_reports.xlsx", 2) %>%
  mutate("Sample" = "NIST-6") %>%
  filter("new_est_reads" > 0)
NIST7  <- read_excel("NIST_data/Bracken_reports.xlsx", 3) %>%
  mutate("Sample" = "NIST-7") %>%
  filter(new_est_reads > 0)
NIST8 <- read_excel("NIST_data/Bracken_reports.xlsx", 4) %>%
  mutate("Sample" = "NIST-8") %>%
  filter(new_est_reads > 0)

NIST_SG <- full_join(NIST5, NIST6, by = c("name", "taxonomy_id", "taxonomy_lvl",
                                          "kraken_assigned_reads", "added_reads", "new_est_reads",
                                          "fraction_total_reads", "Sample")) %>%
  full_join(., NIST7, by = c("name", "taxonomy_id", "taxonomy_lvl",
                                          "kraken_assigned_reads", "added_reads", "new_est_reads",
                                          "fraction_total_reads", "Sample"))%>%
  full_join(., NIST8, by = c("name", "taxonomy_id", "taxonomy_lvl",
                                          "kraken_assigned_reads", "added_reads", "new_est_reads",
                                          "fraction_total_reads", "Sample")) %>%
  rename("BrackenCount" = "new_est_reads") %>%
  rename("KrakenCount" = "kraken_assigned_reads") %>%
  rename("Genus"= "name") %>%
  select(Sample, Genus, BrackenCount, KrakenCount, fraction_total_reads) %>%
  group_by(Sample) %>%
  mutate("TotalBrakenReads" = sum(BrackenCount)) 

#16S data analyzed with Dada2
NIST_16SData <- read_delim("NIST_data/16S_Analysis/dada2_output/taxa_tab.txt",  "\t") %>%
  select(Sample, Genus, Count) %>%
  group_by(Sample, Genus) %>%
  summarise_each(funs(sum)) %>%
  group_by(Sample) %>%
  mutate(Reads = sum(Count)) %>%
  mutate(RelAbund = Count/Reads) %>%
  subset(Sample == "NIST-1" | Sample == "NIST-2"| Sample == "NIST-3"| Sample == "NIST-4") %>%
  mutate_at('Genus', ~replace_na(.,"unclassified"))
```

```
## 
## ── Column specification ────────────────────────────────────────────────────────
## cols(
##   Kingdom = col_character(),
##   Phylum = col_character(),
##   Class = col_character(),
##   Order = col_character(),
##   Family = col_character(),
##   Genus = col_character(),
##   Species = col_character(),
##   Sequence = col_character(),
##   asvID = col_character(),
##   Sample = col_character(),
##   Count = col_double(),
##   RelAbund = col_double()
## )
```

```
#update location if running new analysis
save.image("NIST-analysis.RData")

RA_NIST_16SData <- NIST_16SData %>%
  select(-Count, -Reads) %>%
  mutate("Analysis" = "NIST") %>%
  ungroup() %>%
  select(Sample, Genus, RelAbund, Analysis)
```

import and build Other Donor Table

```
OM1137978 <- read_excel("NIST_data/Bracken_reportsOD.xlsx", 1) %>%
  mutate("Sample" = "1-1") %>%
  filter("new_est_reads" > 0)
OM1137996 <- read_excel("NIST_data/Bracken_reportsOD.xlsx", 2) %>%
  mutate("Sample" = "1-2") %>%
  filter("new_est_reads" > 0)

OM2138056 <- read_excel("NIST_data/Bracken_reportsOD.xlsx", 3) %>%
  mutate("Sample" = "2-1") %>%
  filter(new_est_reads > 0)
OM2138071 <- read_excel("NIST_data/Bracken_reportsOD.xlsx", 4) %>%
  mutate("Sample" = "2-2") %>%
  filter(new_est_reads > 0)

OF1138128 <- read_excel("NIST_data/Bracken_reportsOD.xlsx", 5) %>%
  mutate("Sample" = "3-1") %>%
  filter("new_est_reads" > 0)
OF1138144 <- read_excel("NIST_data/Bracken_reportsOD.xlsx", 6) %>%
  mutate("Sample" = "3-2") %>%
  filter("new_est_reads" > 0)

OF2138209 <- read_excel("NIST_data/Bracken_reportsOD.xlsx", 7) %>%
  mutate("Sample" = "4-1") %>%
  filter("new_est_reads" > 0)
OF2138218 <- read_excel("NIST_data/Bracken_reportsOD.xlsx", 8) %>%
  mutate("Sample" = "4-2") %>%
  filter("new_est_reads" > 0)

VM1139035<- read_excel("NIST_data/Bracken_reportsOD.xlsx", 9) %>%
  mutate("Sample" = "5-1") %>%
  filter("new_est_reads" > 0) 
VM1139043 <- read_excel("NIST_data/Bracken_reportsOD.xlsx", 10) %>%
  mutate("Sample" = "5-2") %>%
  filter("new_est_reads" > 0)

VM2139114 <- read_excel("NIST_data/Bracken_reportsOD.xlsx", 11) %>%
  mutate("Sample" = "6-1") %>%
  filter("new_est_reads" > 0)
VM2139129 <- read_excel("NIST_data/Bracken_reportsOD.xlsx", 12) %>%
  mutate("Sample" = "6-2") %>%
  filter("new_est_reads" > 0)

VF1139189 <- read_excel("NIST_data/Bracken_reportsOD.xlsx", 13) %>%
  mutate("Sample" = "7-1") %>%
  filter("new_est_reads" > 0)
VF1139210 <- read_excel("NIST_data/Bracken_reportsOD.xlsx", 14) %>%
  mutate("Sample" = "7-2") %>%
  filter("new_est_reads" > 0)

VF2139269 <- read_excel("NIST_data/Bracken_reportsOD.xlsx", 15) %>%
  mutate("Sample" = "8-1") %>%
  filter("new_est_reads" > 0)
VF2139288 <- read_excel("NIST_data/Bracken_reportsOD.xlsx", 16) %>%
  mutate("Sample" = "8-2") %>%
  filter("new_est_reads" > 0)

Other_donors <- full_join(OM1137978, OM1137996, by = c("name", "taxonomy_id", "taxonomy_lvl",
                                                       "kraken_assigned_reads", "added_reads",
                                                       "new_est_reads", "fraction_total_reads",
                                                       "Sample")) %>%
  full_join(., OM2138056, by = c("name", "taxonomy_id", "taxonomy_lvl",
                                                       "kraken_assigned_reads", "added_reads",
                                                       "new_est_reads", "fraction_total_reads",
                                                       "Sample")) %>%
  full_join(., OM2138071, by = c("name", "taxonomy_id", "taxonomy_lvl",
                                                       "kraken_assigned_reads", "added_reads",
                                                       "new_est_reads", "fraction_total_reads",
                                                       "Sample")) %>%
  full_join(., OF1138128, by = c("name", "taxonomy_id", "taxonomy_lvl",
                                                       "kraken_assigned_reads", "added_reads",
                                                       "new_est_reads", "fraction_total_reads",
                                                       "Sample")) %>%
  full_join(., OF1138144, by = c("name", "taxonomy_id", "taxonomy_lvl",
                                                       "kraken_assigned_reads", "added_reads",
                                                       "new_est_reads", "fraction_total_reads",
                                                       "Sample")) %>%
  full_join(., OF2138209, by = c("name", "taxonomy_id", "taxonomy_lvl",
                                                       "kraken_assigned_reads", "added_reads",
                                                       "new_est_reads", "fraction_total_reads",
                                                       "Sample")) %>%
  full_join(., OF2138218, by = c("name", "taxonomy_id", "taxonomy_lvl",
                                                       "kraken_assigned_reads", "added_reads",
                                                       "new_est_reads", "fraction_total_reads",
                                                       "Sample")) %>%
  full_join(., VM1139035, by = c("name", "taxonomy_id", "taxonomy_lvl",
                                                       "kraken_assigned_reads", "added_reads",
                                                       "new_est_reads", "fraction_total_reads",
                                                       "Sample")) %>%
  full_join(., VM1139043, by = c("name", "taxonomy_id", "taxonomy_lvl",
                                                       "kraken_assigned_reads", "added_reads",
                                                       "new_est_reads", "fraction_total_reads",
                                                       "Sample")) %>%
  full_join(., VM2139114, by = c("name", "taxonomy_id", "taxonomy_lvl",
                                                       "kraken_assigned_reads", "added_reads",
                                                       "new_est_reads", "fraction_total_reads",
                                                       "Sample")) %>%
  full_join(., VM2139129, by = c("name", "taxonomy_id", "taxonomy_lvl",
                                                       "kraken_assigned_reads", "added_reads",
                                                       "new_est_reads", "fraction_total_reads",
                                                       "Sample")) %>%
  full_join(., VF1139189, by = c("name", "taxonomy_id", "taxonomy_lvl",
                                                       "kraken_assigned_reads", "added_reads",
                                                       "new_est_reads", "fraction_total_reads",
                                                       "Sample")) %>%
  full_join(., VF1139210, by = c("name", "taxonomy_id", "taxonomy_lvl",
                                                       "kraken_assigned_reads", "added_reads",
                                                       "new_est_reads", "fraction_total_reads",
                                                       "Sample")) %>%
  full_join(., VF2139269, by = c("name", "taxonomy_id", "taxonomy_lvl",
                                                       "kraken_assigned_reads", "added_reads",
                                                       "new_est_reads", "fraction_total_reads",
                                                       "Sample")) %>%
  full_join(., VF2139288, by = c("name", "taxonomy_id", "taxonomy_lvl",
                                                       "kraken_assigned_reads", "added_reads",
                                                       "new_est_reads", "fraction_total_reads",
                                                       "Sample")) %>%
  rename("BrackenCount" = "new_est_reads") %>%
  rename("KrakenCount" = "kraken_assigned_reads") %>%
  rename("Genus"= "name") %>%
  select(Sample, Genus, BrackenCount, KrakenCount, fraction_total_reads) %>%
  group_by(Sample) %>%
  mutate("TotalBrakenReads" = sum(BrackenCount))
```

```
Genus_NISTData <- full_join(NIST_SG, Other_donors, by = c("Sample", "Genus", "BrackenCount", "KrakenCount",
                                                          "fraction_total_reads","TotalBrakenReads")) 

GenusRead_count <- Genus_NISTData %>%
  select(Sample, TotalBrakenReads) %>%
  distinct(Sample, TotalBrakenReads)

RA_NIST_SGData <- Genus_NISTData %>%
  select(Sample, Genus, fraction_total_reads) %>%
  rename("RelAbund" = "fraction_total_reads") %>%
  mutate("Analysis" = "NIST")

#update location if running new analysis
save.image("NIST-analysis.RData")
```

### Select single donor data from NIST data set, same donor that was sent to all DTC companies - no other biological samples

```
Donor5_NISTData <- NIST_SG %>%
  select(Sample, Genus, fraction_total_reads) %>%
  rename("RelAbund" = "fraction_total_reads") %>%
  mutate("Analysis" = "NIST") %>%
  bind_rows(RA_NIST_16SData) 

RA_Donor5Data <- Donor5_NISTData %>%
  bind_rows(RA_DTCData) %>%
  unite(Sample, Sample, Analysis, sep = "_", remove = T)

Donor5.Genus.Table <- RA_Donor5Data %>%
  spread(key = Genus, value = RelAbund) %>%
  replace(is.na(.), 0)

Donor5_allunique <- RA_Donor5Data %>%
  distinct(Genus, .keep_all = T)
```

### combining all Data into single Genus Table formated for diversity tests

```
RA_fullData <- bind_rows(RA_DTCData, RA_NIST_16SData) %>%
  bind_rows(RA_NIST_SGData) %>%
  unite(Sample, Sample, Analysis, sep = "_", remove = T) %>%
  mutate(Genus = str_replace(Genus,"unclassified", "Unspecified"))

group_by(RA_fullData, Sample) %>%
  summarise(sum(RelAbund))
```

```
## # A tibble: 44 x 2
##    Sample   `sum(RelAbund)`
##    <chr>              <dbl>
##  1 1-1_NIST            1.00
##  2 1-2_NIST            1.00
##  3 2-1_NIST            1.00
##  4 2-2_NIST            1.00
##  5 3-1_NIST            1.00
##  6 3-2_NIST            1.00
##  7 4-1_NIST            1.00
##  8 4-2_NIST            1   
##  9 5-1_NIST            1.00
## 10 5-2_NIST            1.00
## # … with 34 more rows
```

```
SampleOrder <- c("A-1_DTC","A-2_DTC","A-3_DTC","B-1_DTC","B-2_DTC","B-3_DTC", "C-1_DTC","C-2_DTC","C-3_DTC",
                 "D-1_DTC","D-2_DTC","D-3_DTC","E-1_DTC","E-2_DTC","E-3_DTC", "F-1_DTC","F-2_DTC","F-3_DTC",
                 "G-1_DTC","G-2_DTC","NIST-1_NIST","NIST-2_NIST","NIST-3_NIST", "NIST-4_NIST", 
                 "NIST-5_NIST", "NIST-6_NIST", "NIST-7_NIST", "NIST-8_NIST", 
                 "1-1_NIST", "1-2_NIST", "2-1_NIST", "2-2_NIST", "3-1_NIST", "3-2_NIST", "4-1_NIST", "4-2_NIST",
                 "5-1_NIST", "5-2_NIST","6-1_NIST", "6-2_NIST","7-1_NIST", "7-2_NIST", "8-1_NIST", "8-2_NIST")

#reformate data frame into wide version Sample as rows, Genera as columns, RA for value
Full.Genus.Table <- RA_fullData %>%
  spread(key = Genus, value = RelAbund) %>%
  replace(is.na(.), 0) %>%
  arrange(factor(Sample, levels = SampleOrder))

save.image("NIST-analysis.RData")
```

### Calculate alpha diversity metrics

```
InvSimpson <- Full.Genus.Table %>%
  column_to_rownames(var="Sample") %>%
  diversity(., MARGIN = 1, index = "invsimpson") %>%
  as.data.frame() %>%
  rownames_to_column("Sample") %>%
  dplyr::rename("InvSimpson" = ".")

Shannon <- Full.Genus.Table %>%
  column_to_rownames(var="Sample") %>%
  diversity(., MARGIN = 1, index = "shannon") %>%
  as.data.frame() %>%
  rownames_to_column("Sample") %>%
  dplyr::rename("Shannon" = ".")

sppr <- Full.Genus.Table %>%
  column_to_rownames(var="Sample") %>%
  specnumber() %>%
  as.data.frame() %>%
  rownames_to_column("Sample") %>%
  dplyr::rename("Genus_count" = ".")

alphadiversity <- full_join(Shannon, InvSimpson, by = "Sample") %>%
  full_join(., sppr, by = "Sample") %>%
  separate(Sample, sep = "_", into=c("Sample","Analysis")) %>%
  full_join(Meta, by = "Sample") 

save.image("NIST-analysis.RData")
```

### generate alpha Diversity Figures

```
InvSimpson_plot <- alphadiversity %>%
  filter(Company != "I") %>%
  filter(Donor == "Donor5") %>%
  ggplot(., aes(x = Company, y = InvSimpson)) + 
  geom_point(size = 3, stroke = 1.2, shape = 0) +
    labs(x = "Company",
       y = "Inverse Simpson") +
  theme_gray() +
  theme(text = element_text(size = 20), axis.text.x = element_text(angle=45, vjust=1, hjust=1)) +
  facet_wrap(.~MGS, scales = "free_x")

print(InvSimpson_plot)
```

```
Shannon_plot <- alphadiversity %>%
  filter(Company != "I") %>%
  filter(Donor == "Donor5") %>%
  ggplot(., aes(x = Company, y = Shannon)) + 
  geom_point(size = 3, stroke = 1.2, shape = 1) +
    labs(x = "Company",
       y = "Shannon Diversity") +
  theme_gray() +
  theme(text = element_text(size = 20), axis.text.x = element_text(angle=45, vjust=1, hjust=1)) +
  facet_wrap(.~MGS, scales = "free_x")

print(Shannon_plot)
```

```
GenusCount_plot <- alphadiversity %>%
  filter(Company != "I") %>%
  filter(Donor == "Donor5") %>%
  ggplot(., aes(x = Company, y = Genus_count)) + 
  geom_point(size = 3, stroke = 1.2, shape = 2) +
  theme_gray() +
    labs(x = "Company",
       y = "Genera identified")+
  theme(text = element_text(size = 20), axis.text.x = element_text(angle=45, vjust=1, hjust=1)) +
  facet_wrap(.~MGS, scales = "free_x")
print(GenusCount_plot)
```

```
ggsave("saved_plots/InvSimpson.png",InvSimpson_plot, width = 4, height = 4, units = "in")
ggsave("saved_plots/Shannon.png",Shannon_plot, width = 4, height = 4, units = "in")
ggsave("saved_plots/Genus_count.png",GenusCount_plot, width = 4, height = 4, units = "in")
```

```
beta_dist <- Full.Genus.Table %>%
  column_to_rownames(var="Sample") %>%
  vegdist(., index = "bray", k = 2) 

mds <- metaMDS(beta_dist)
```

```
## Run 0 stress 0.1416492 
## Run 1 stress 0.143486 
## Run 2 stress 0.1367111 
## ... New best solution
## ... Procrustes: rmse 0.02843059  max resid 0.1727015 
## Run 3 stress 0.1416489 
## Run 4 stress 0.1367111 
## ... Procrustes: rmse 0.0002534321  max resid 0.001307568 
## ... Similar to previous best
## Run 5 stress 0.1367111 
## ... New best solution
## ... Procrustes: rmse 5.51084e-05  max resid 0.0002679689 
## ... Similar to previous best
## Run 6 stress 0.136711 
## ... New best solution
## ... Procrustes: rmse 2.7272e-05  max resid 0.0001170499 
## ... Similar to previous best
## Run 7 stress 0.1416489 
## Run 8 stress 0.1367111 
## ... Procrustes: rmse 3.294773e-05  max resid 0.0001320231 
## ... Similar to previous best
## Run 9 stress 0.136711 
## ... New best solution
## ... Procrustes: rmse 0.0001008157  max resid 0.0005449314 
## ... Similar to previous best
## Run 10 stress 0.1527926 
## Run 11 stress 0.1367111 
## ... Procrustes: rmse 2.114146e-05  max resid 8.870964e-05 
## ... Similar to previous best
## Run 12 stress 0.136711 
## ... New best solution
## ... Procrustes: rmse 7.35988e-05  max resid 0.0004176627 
## ... Similar to previous best
## Run 13 stress 0.1414668 
## Run 14 stress 0.141649 
## Run 15 stress 0.1416489 
## Run 16 stress 0.1559879 
## Run 17 stress 0.136711 
## ... Procrustes: rmse 4.747477e-06  max resid 1.963755e-05 
## ... Similar to previous best
## Run 18 stress 0.1414668 
## Run 19 stress 0.141467 
## Run 20 stress 0.1367111 
## ... Procrustes: rmse 6.110106e-05  max resid 0.0003014164 
## ... Similar to previous best
## *** Best solution repeated 3 times
```

```
mds_data <- as.data.frame(mds$points) %>%
  rownames_to_column("Sample") %>%
  separate(Sample, sep = "_", into=c("Sample","Analysis")) %>%
  full_join(Meta, by = "Sample") 

stressplot(mds)
```

```
plot(mds)
```

```
## species scores not available
```

```
mds_data2 <- mds_data %>%
  mutate(Group1 = ifelse(Donor == "Donor5", "Donor5", "Other")) %>%
  unite(Group2, c("Group1", "MGS"), remove = F)
```

```
beta_dist_donor5 <- Donor5.Genus.Table %>%
  column_to_rownames(var="Sample") %>%
  vegdist(., index = "bray", k = 2) 

mds_donor5 <- metaMDS(beta_dist_donor5)
```

```
## Run 0 stress 0.09815314 
## Run 1 stress 0.09815316 
## ... Procrustes: rmse 0.0001960264  max resid 0.0005448906 
## ... Similar to previous best
## Run 2 stress 0.0981533 
## ... Procrustes: rmse 0.0002774357  max resid 0.0007864819 
## ... Similar to previous best
## Run 3 stress 0.09815311 
## ... New best solution
## ... Procrustes: rmse 5.664913e-05  max resid 0.0001505572 
## ... Similar to previous best
## Run 4 stress 0.1013888 
## Run 5 stress 0.1013887 
## Run 6 stress 0.1007051 
## Run 7 stress 0.1013889 
## Run 8 stress 0.1007053 
## Run 9 stress 0.1013889 
## Run 10 stress 0.1013889 
## Run 11 stress 0.1013889 
## Run 12 stress 0.09815309 
## ... New best solution
## ... Procrustes: rmse 1.313676e-05  max resid 4.639266e-05 
## ... Similar to previous best
## Run 13 stress 0.09815319 
## ... Procrustes: rmse 0.0001532933  max resid 0.0004243178 
## ... Similar to previous best
## Run 14 stress 0.1013889 
## Run 15 stress 0.1007019 
## Run 16 stress 0.09815316 
## ... Procrustes: rmse 9.10788e-05  max resid 0.0002796147 
## ... Similar to previous best
## Run 17 stress 0.1007051 
## Run 18 stress 0.09815328 
## ... Procrustes: rmse 0.000205355  max resid 0.000567461 
## ... Similar to previous best
## Run 19 stress 0.1013889 
## Run 20 stress 0.1007018 
## *** Best solution repeated 4 times
```

```
mds_data_donor5 <- as.data.frame(mds_donor5$points) %>%
  rownames_to_column("Sample") %>%
  separate(Sample, sep = "_", into=c("Sample","Analysis")) %>%
  inner_join(Meta)
```

```
## Joining, by = "Sample"
```

```
stressplot(mds_donor5)
```

```
plot(mds_donor5)
```

```
## species scores not available
```

```
BetaDiv5 <- mds_data_donor5 %>%
  ggplot() +
  geom_point(aes(x = MDS1, y = MDS2, color = Company, shape = Donor), size = 5, alpha = 1, stroke = 1) +
  geom_mark_ellipse(aes(x = MDS1, y = MDS2, fill = Company, color = Company, linetype = MGS),alpha = 0.1) +
  theme_bw() +
  theme(text = element_text(size = 15)) +
  scale_shape_manual(values = 8) +
  scale_color_manual(values = pal_sample) +
  scale_fill_manual(values = pal_sample) +
  scale_linetype_manual(values = c("longdash","solid")) +
  scale_x_continuous(limits = c(-2,1)) +
  scale_y_continuous(limits = c(-1, 1)) +
  geom_vline(xintercept = c(0), color = "grey70", linetype = 2) +
  geom_hline(yintercept = c(0), color = "grey70", linetype = 2) +
  labs(title = "Donor5-Beta-diversity") 

print(BetaDiv5)
```

```
BetaDivAll <- ggplot(mds_data2) +
  geom_point(aes(x = MDS1, y = MDS2, shape = Donor), size = 5, alpha = 1, stroke = 1) +
  geom_mark_ellipse(aes(x = MDS1, y = MDS2, fill = Group1, linetype = MGS), alpha = 0.1) +
  theme_bw() +
  theme(text = element_text(size = 15)) +
  scale_shape_manual(values=c(24, 21, 22, 23, 8, 15, 17, 18)) +
  scale_color_manual(values = c("black", "blue", "darkgreen")) +
  scale_fill_manual(values = c( "blue", "yellow")) +
   scale_linetype_manual(values = c("longdash","solid")) +
  scale_x_continuous(limits = c(-2,1)) +
  scale_y_continuous(limits = c(-1, 1)) +
  geom_vline(xintercept = c(0), color = "grey70", linetype = 2) +
  geom_hline(yintercept = c(0), color = "grey70", linetype = 2) +
  labs(title = "FullCohort-Beta-diversity") 

print(BetaDivAll)
```

```
ggsave("saved_plots/betadiv_plot1.png", BetaDiv5, width = 7, height = 5, units = "in")
ggsave("saved_plots/betadiv_plot2.png", BetaDivAll, width = 7, height = 5, units = "in")

save.image("NIST-analysis.RData")
```

### common genera only

### semi join returns only rows in X that have a match in y; semi\_join = (x,y, by = Genus)

```
#selecting genera in common between NIST 16S data sets
NIST1_16S <- NIST_16SData %>%
  subset(Sample == "NIST-1") %>%
  filter(Count != 0) %>%
  select(-Count, -Reads) %>%
  spread(key = "Sample", value = "RelAbund")

NIST2_16S <- NIST_16SData %>%
  subset(Sample == "NIST-2") %>%
  filter(Count != 0)%>%
  select(-Count, -Reads) %>%
  spread(key = "Sample", value = "RelAbund")

NIST3_16S <- NIST_16SData %>%
  subset(Sample == "NIST-3") %>%
  filter(Count != 0)%>%
  select(-Count, -Reads) %>%
  spread(key = "Sample", value = "RelAbund")

NIST4_16S <- NIST_16SData %>%
  subset(Sample == "NIST-4") %>%
  filter(Count != 0)%>%
  select(-Count, -Reads) %>%
  spread(key = "Sample", value = "RelAbund")

common_NIST16S_genera <- inner_join(NIST1_16S, NIST2_16S, by = "Genus") %>%
  inner_join(NIST3_16S, by = "Genus") %>%
  inner_join(NIST4_16S, by = "Genus") 

#common NIST SG genera
NIST5_SG <- NIST_SG %>%
  subset(Sample == "NIST-5") %>%
  select(Sample, Genus, fraction_total_reads) %>%
  dplyr::rename("RelAbund" = "fraction_total_reads") %>%
  spread(key = "Sample", value = "RelAbund")

NIST6_SG <- NIST_SG %>%
  subset(Sample == "NIST-6") %>%
  select(Sample, Genus, fraction_total_reads) %>%
  dplyr::rename("RelAbund" = "fraction_total_reads") %>%
  spread(key = "Sample", value = "RelAbund")

NIST7_SG <- NIST_SG %>%
  subset(Sample == "NIST-7") %>%
  select(Sample, Genus, fraction_total_reads) %>%
  dplyr::rename("RelAbund" = "fraction_total_reads") %>%
  spread(key = "Sample", value = "RelAbund")

NIST8_SG <- NIST_SG %>%
  subset(Sample == "NIST-8")%>%
  select(Sample, Genus, fraction_total_reads) %>%
  dplyr::rename("RelAbund" = "fraction_total_reads") %>%
  spread(key = "Sample", value = "RelAbund")

common_NISTSG_genera <- inner_join(NIST5_SG, NIST6_SG, by = "Genus")%>%
  inner_join(., NIST7_SG, by = "Genus") %>%
  inner_join(., NIST8_SG, by = "Genus") 

#common NIST Genera
common_NIST_genera <- inner_join (common_NIST16S_genera, common_NISTSG_genera, by = "Genus") 

#Genera common to DTC
common_DTC_generaALL <- DTC.Genus.table %>%
  filter(Genus != "Unspecified") %>%
  na.omit()

common_DTC_genera <- select(DTC.Genus.table, -"A-3") %>%
  na.omit()
  
#common to all
common_genera <- inner_join(common_NIST_genera, common_DTC_genera, by = "Genus")
```

```
Unspecified_RA <- filter(RA_fullData, Genus == "Unspecified") %>%
  separate(Sample, sep = "_", into=c("Sample","Analysis")) %>%
  select(-Analysis) %>%
  spread(key = Sample, value = RelAbund) 

Meta5 <- subset(Meta, Donor == "Donor5") %>%
  mutate(Rep = c(1,2,3,1,2,3,1,2,3,1,2,3,4,
                 1,2,3,1,2,3,1,2,3,1,2,5,6,7,8,9,10)) %>%
  unite(ID, c(Company, Rep), sep= "-", remove = F)

print(Unspecified_RA)
```

```
## # A tibble: 1 x 25
##   Genus     `A-1` `A-2` `A-3` `B-1` `B-2` `B-3`  `C-1`  `C-2`  `C-3` `D-1` `D-2`
##   <chr>     <dbl> <dbl> <dbl> <dbl> <dbl> <dbl>  <dbl>  <dbl>  <dbl> <dbl> <dbl>
## 1 Unspecif… 0.238 0.207 0.553 0.474 0.467 0.453 0.0799 0.0281 0.0275     0     0
## # … with 13 more variables: D-3 <dbl>, E-1 <dbl>, E-2 <dbl>, E-3 <dbl>,
## #   F-1 <dbl>, F-2 <dbl>, F-3 <dbl>, G-1 <dbl>, G-2 <dbl>, NIST-1 <dbl>,
## #   NIST-2 <dbl>, NIST-3 <dbl>, NIST-4 <dbl>
```

```
common_genera_plot <- common_genera %>%
  rbind.fill(Unspecified_RA) %>%
  replace(is.na(.), 0) %>%
  select(-"A-3") %>%
  gather(key = Sample, value = RelAbund, -Genus) %>%
  inner_join(., Meta5, by = "Sample") %>%
  ggplot(aes(x=ID, y=RelAbund, fill = Genus)) +
  geom_bar(stat = "identity", show.legend = T) +
  scale_fill_manual(values = pal_35) +
  scale_y_continuous(labels = scales::percent) +
  theme_linedraw() +
  theme(axis.text.y = element_text(size = 15), axis.text.x=element_text(angle=45,hjust=1,size=10),
        axis.title = element_text(size = 20), legend.text = element_text(size = 12)) 

print(common_genera_plot)
```

```
ggsave("saved_plots/common_genera.png",common_genera_plot, width = 11.5, height = 8, units = "in")  

save.image("NIST-analysis.RData")
```

### note: NIST SG data comes from a Bracken Report (H-5 to H-8), during the Bracken analysis unclassified reads are dropped, everything remaining was classified to atleast the genus level. Some companies may have something similar (for example company D has 0 Unspecified reads which could be because the unclassified are dropped from the final calculations)

```
SelectTaxa.Genus.Table <- Full.Genus.Table %>%
  separate(Sample, sep = "_", into=c("Sample","Analysis")) %>%
  full_join(Meta, by = "Sample") %>%
  select(Sample, Analysis, Company, MGS, Donor, Akkermansia, Bacteroides, Bifidobacterium, Clostridium, Enterococcus, Escherichia, Faecalibacterium, Klebsiella,
         Lactobacillus, Prevotella, Roseburia, Ruminococcus, Company) %>%
  gather(key = "Genus", value = "RelAbund", -Sample, -Analysis, -MGS, -Company, -Donor) %>%
  mutate(RelAbund_Perc = RelAbund*100) %>%
  filter(Company != "NIST") %>%
  select(-RelAbund) %>%
  spread(key = Genus, value = RelAbund_Perc)
```

```
Clostridium <- SelectTaxa.Genus.Table %>%
  filter(Company != "I") %>%
  filter(Company != "H") %>%
  ggplot(aes(x=Company, y=Clostridium, shape = MGS)) +
  geom_point(size = 3.5, stroke = 1, color = "blue4",show.legend = F) +
  scale_shape_manual(values = c(1,4)) +
  geom_hline(yintercept=0.06, linetype="dotted", color = "gold") +
  geom_hline(yintercept=0.28, linetype="dotted", color = "gold") +
  geom_hline(yintercept=1.17, linetype="dashed", color = "grey37") +
  geom_rect(xmin = -Inf, xmax = Inf, ymin=0.06, ymax=0.28, alpha=0.01, fill='gold') +
  ylab("RelAbund(%)") +
  ggtitle("Clostridium") +
  theme_linedraw() 

print(Clostridium)
```

```
Bacteroides <- SelectTaxa.Genus.Table %>%
  filter(Company != "I") %>%
  filter(Company != "H") %>%
  ggplot(aes(x=Company, y=Bacteroides, shape = MGS)) +
  geom_point(size = 3.5, stroke = 1, color = "blue4",show.legend = F) +
  scale_shape_manual(values = c(1,4)) +
  geom_hline(yintercept=6.17, linetype="dotted", color = "gold") +
  geom_hline(yintercept=16.73, linetype="dotted", color = "gold") +
  geom_hline(yintercept=29.09, linetype="dashed", color = "grey37") +
  geom_rect(xmin = -Inf, xmax = Inf, ymin=6.17, ymax=16.73, alpha=0.01, fill='gold') +
  ylab("RelAbund(%)") +
  ggtitle("Bacteroides") +
  theme_linedraw()

print(Bacteroides)
```

```
Bifidobacterium <- SelectTaxa.Genus.Table %>%
  filter(Company != "I") %>%
  filter(Company != "H") %>%
  ggplot(aes(x=Company, y=Bifidobacterium, shape = MGS)) +
  geom_point(size = 3.5, stroke = 1, color = "blue4",show.legend = F) +
  scale_shape_manual(values = c(1,4)) +
  geom_hline(yintercept=0.24, linetype="dotted", color = "gold") +
  geom_hline(yintercept=2.82, linetype="dotted", color = "gold") +
  geom_hline(yintercept=1.13, linetype="dashed", color = "grey37") +
  geom_rect(xmin = -Inf, xmax = Inf, ymin=0.24, ymax=2.82, alpha=0.01, fill='gold') +
  ylab("RelAbund(%)") +
  ggtitle("Bifidobacterium") +
  theme_linedraw() 

print(Bifidobacterium)
```

```
Roseburia <- SelectTaxa.Genus.Table %>%
  filter(Company != "I") %>%
  filter(Company != "H") %>%
  ggplot(aes(x=Company, y=Roseburia, shape = MGS)) +
  geom_point(size = 3.5, stroke = 1, color = "blue4",show.legend = F) +
  scale_shape_manual(values = c(1,4)) +
  geom_hline(yintercept=0.73, linetype="dotted", color = "gold") +
  geom_hline(yintercept=3.49, linetype="dotted", color = "gold") +
  geom_hline(yintercept=3.91, linetype="dashed", color = "grey37") +
  geom_rect(xmin = -Inf, xmax = Inf, ymin=0.73, ymax=3.49, alpha=0.01, fill='gold') +
  ylab("RelAbund(%)") +
  ggtitle("Roseburia") +
  theme_linedraw() 

print(Roseburia)
```

```
Faecalibacterium <- SelectTaxa.Genus.Table %>%
  filter(Company != "I") %>%
  filter(Company != "H") %>%
  ggplot(aes(x=Company, y=Faecalibacterium, shape = MGS)) +
  geom_point(size = 3.5, stroke = 1,color = "blue4", show.legend = F) +
  scale_shape_manual(values = c(1,4)) +
  geom_hline(yintercept=0.64, linetype="dotted", color = "gold") +
  geom_hline(yintercept=2.01, linetype="dotted", color = "gold") +
  geom_hline(yintercept=8.73, linetype="longdash", color = "grey37") +
  geom_rect(xmin = -Inf, xmax = Inf, ymin=0.64, ymax=2.01, alpha=0.01, fill='gold') +
  ylab("RelAbund(%)") +
  ggtitle("Faecalibacterium") +
  theme_linedraw() 

print(Faecalibacterium)
```

```
ggsave("saved_plots/Clostridium.png", Clostridium, width = 5, height = 3, units = "in")
ggsave("saved_plots/Bacteroides.png", Bacteroides, width = 5, height = 3, units = "in")
ggsave("saved_plots/Bifidobacterium.png", Bifidobacterium, width = 5, height = 3, units = "in")
ggsave("saved_plots/Roseburia.png", Roseburia, width = 5, height = 3, units = "in")  
ggsave("saved_plots/Faecalibacterium.png", Faecalibacterium, width = 5, height = 3, units = "in")
save.image("NIST-analysis.RData")
```

### Comparison of reproducibility of Company A replicates

```
DTC_CompanyA <- select(DTC.Genus.table, Genus, "A-1", "A-2", "A-3") %>%
  gather(key = Sample, value = RelAbund, -Genus) %>%
  inner_join(Meta, by = "Sample") %>%
  na.omit()
  
CompanyA_common <- select(DTC.Genus.table, Genus, "A-1", "A-2", "A-3") %>%
  na.omit() %>%
  gather(key = Sample, value = RelAbund, -Genus)
```

```
A3_unique <- DTC_CompanyA %>%
  filter(Sample == "A-3")

genus_sumA3 <- aggregate(RelAbund ~ Genus, data = A3_unique, FUN = sum)
genus_sumA3 <- genus_sumA3[order(genus_sumA3$RelAbund), ]

A3_unique_plot <- A3_unique %>%
  ggplot(aes(x=Sample, y=RelAbund, fill = factor(Genus, levels = genus_sumA3$Genus))) +
  geom_bar(stat = "identity", color = "white", size = 0.01, show.legend = T) +
  scale_fill_manual(values = c("Abiotrophia" = "#1F78C8" ,"Actinomyces" = "#ff0000","Aggregatibacter" = "#33a02c","Alloprevotella" = "#6A33C2","Anaeroglobus"= "#ff7f00","Atopobium"= "#FFD700",
                               "Campylobacter"= "#a6cee3","Capnocytophaga"= "#FB6496","Cardiobacterium"= "#808080", "Centipeda" = "#CAB2D6","Corynebacterium"= "#b2df8a","Dialister"= "#191919",
                               "Eubacterium" = "#EEE685","Fusobacterium" = "#C8308C","Gemella" = "#FF83FA", "Granulicatella" = "#C814FA","Haemophilus" = "#0000FF",
                               "Lachnoanaerobaculum" = "#36648B",
                               "Lautropia"= "#565656","Leptotrichia"= "#00E2E5","Mogibacterium"= "#778B00","Neisseria"= "#BEBE00","Oribacterium"= "#8B3B00", "Parvimonas"= "#00FF00", 
                               "Peptostreptococcus"= "#A52A3C", "Prevotella"= "#F0A0FF", "Pseudopropionibacterium"= "#0075DC", "Rothia"= "#993F00","Scardovia"= "#4C005C", "Slackia"= "#005C31", 
                               "Solobacterium"= "#2BCE48", "Stomatobaculum"= "#FFCC99", "Streptococcus"= "#FDBF6F", "Veillonella"= "#94FFB5", "Unspecified" = "#999999")) +
 
  labs(fill = "Genus") +
  scale_y_continuous(labels = scales::percent) +
  theme(axis.text.x=element_text(angle=90,hjust=1)) +
  theme(axis.text.y = element_text(size = 15), axis.text.x=element_text(angle=90,hjust=1,size=15),
        axis.title = element_text(size = 20), legend.text = element_text(size = 12)) +
  theme_linedraw()
```

```
## Warning: Using `size` aesthetic for lines was deprecated in ggplot2 3.4.0.

## Warning: Please use `linewidth` instead.
```

```
print(A3_unique_plot)
```

```
genus_sum <- aggregate(RelAbund ~ Genus, data = CompanyA_common, FUN = sum)
genus_sum <- genus_sum[order(genus_sum$RelAbund), ]

CompanyA_common_plot <- CompanyA_common %>%
  ggplot(aes(x=Sample, y=RelAbund, fill = factor(Genus, levels = genus_sum$Genus))) +
  geom_bar(stat = "identity", color = "white", size = 0.01, show.legend = T) +
  scale_fill_manual(values = c("Actinomyces" = "#ff0000","Veillonella"= "#94FFB5","Haemophilus" = "#0000FF", "Dialister"= "#191919",
                               "Streptococcus"="#FDBF6F","Eubacterium" = "#EEE685","Unspecified" = "#999999")) +
  scale_y_continuous(limits = c(0, 0.8), labels = scales::percent) +
  theme(axis.text.x=element_text(angle=90,hjust=1)) +
  theme(axis.text.y = element_text(size = 15), axis.text.x=element_text(angle=90,hjust=1,size=15),
        axis.title = element_text(size = 20), legend.text = element_text(size = 12)) +
  labs(fill = "Genus") +
  theme_linedraw()

print(CompanyA_common_plot)
```

```
ggsave("saved_plots/A_unique1.png", A3_unique_plot, width = 7, height = 5, units = "in")
ggsave("saved_plots/CompanyA_common.png",CompanyA_common_plot, width = 5, height = 5, units = "in")
save.image("NIST-analysis.RData")
```

### how reproducible were results between replicates (for comparison to company A)

```
A_CommonG  <- inner_join(Genus.A1, Genus.A2, by = "Genus") %>%
  inner_join(., Genus.A3, by = "Genus") %>%
  filter(Genus != "Unspecified")

B_CommonG <- inner_join(Genus.B2, Genus.B3, by = "Genus") %>%
  inner_join(., Genus.B1, by = "Genus") %>%
  filter(Genus != "Unspecified")

C_CommonG <- inner_join(Genus.C1, Genus.C2, by = "Genus") %>%
  inner_join(., Genus.C3, by = "Genus") %>%
  filter(Genus != "Unspecified")

D_CommonG <- inner_join(Genus.D1, Genus.D2, by = "Genus") %>%
  inner_join(., Genus.D3, by = "Genus") %>%
  filter(Genus != "Unspecified")

E_CommonG <- inner_join(Genus.E1, Genus.E2, by = "Genus") %>%
  inner_join(., Genus.E3, by = "Genus") %>%
  filter(Genus != "Unspecified")

F_CommonG <- inner_join(Genus.F1, Genus.F2, by = "Genus") %>%
  inner_join(., Genus.F3, by = "Genus") %>%
  filter(Genus != "Unspecified")

G_CommonG <- inner_join(Genus.G1, Genus.G2, by = "Genus") %>%
  filter(Genus != "unclassified")

NIST16S_CommonG <- inner_join(NIST1_16S, NIST2_16S, by = "Genus") %>%
  inner_join(., NIST3_16S, by = "Genus") %>%
  inner_join(., NIST4_16S, by = "Genus") %>%
  filter(Genus != "Unspecified")

NISTSG_CommonG <- inner_join(NIST5_SG, NIST6_SG, by = "Genus") %>%
  inner_join(., NIST7_SG, by = "Genus") %>%
  inner_join(., NIST8_SG, by = "Genus") %>%
  filter(Genus != "Unspecified")

#determine % of sample classified and conserved
A_CommonG  %>%
  summarise_if(is.numeric, sum, na.rm = TRUE)
```

```
## # A tibble: 1 x 3
##   `A-1` `A-2` `A-3`
##   <dbl> <dbl> <dbl>
## 1 0.105 0.106 0.163
```

```
B_CommonG  %>%
  summarise_if(is.numeric, sum, na.rm = TRUE)
```

```
## # A tibble: 1 x 3
##   `B-2` `B-3` `B-1`
##   <dbl> <dbl> <dbl>
## 1 0.523 0.538 0.524
```

```
C_CommonG  %>%
  summarise_if(is.numeric, sum, na.rm = TRUE)
```

```
## # A tibble: 1 x 3
##   `C-1` `C-2` `C-3`
##   <dbl> <dbl> <dbl>
## 1 0.917 0.967 0.969
```

```
D_CommonG  %>%
  summarise_if(is.numeric, sum, na.rm = TRUE)
```

```
## # A tibble: 1 x 3
##   `D-1` `D-2` `D-3`
##   <dbl> <dbl> <dbl>
## 1  1.02 0.996 0.997
```

```
E_CommonG  %>%
  summarise_if(is.numeric, sum, na.rm = TRUE)
```

```
## # A tibble: 1 x 3
##   `E-1` `E-2` `E-3`
##   <dbl> <dbl> <dbl>
## 1 0.857 0.789 0.841
```

```
F_CommonG  %>%
  summarise_if(is.numeric, sum, na.rm = TRUE)
```

```
## # A tibble: 1 x 3
##   `F-1` `F-2` `F-3`
##   <dbl> <dbl> <dbl>
## 1 0.295 0.293 0.296
```

```
G_CommonG %>%
  summarise_if(is.numeric, sum, na.rm = TRUE)
```

```
## # A tibble: 1 x 2
##   `G-1` `G-2`
##   <dbl> <dbl>
## 1 0.999 0.999
```

```
NIST16S_CommonG %>%
  summarise_if(is.numeric, sum, na.rm = TRUE)
```

```
## # A tibble: 1 x 4
##   `NIST-1` `NIST-2` `NIST-3` `NIST-4`
##      <dbl>    <dbl>    <dbl>    <dbl>
## 1    0.998    0.999    0.999    0.998
```

```
NISTSG_CommonG %>%
  summarise_if(is.numeric, sum, na.rm = TRUE)
```

```
## # A tibble: 1 x 4
##   `NIST-5` `NIST-6` `NIST-7` `NIST-8`
##      <dbl>    <dbl>    <dbl>    <dbl>
## 1    0.997    0.996    0.993    0.993
```

```
#determine % of sample classified 
Genus_A %>%
  filter(Genus != "Unspecified") %>%
  summarise_if(is.numeric, sum, na.rm = TRUE)
```

```
## # A tibble: 1 x 3
##   `A-1` `A-2` `A-3`
##   <dbl> <dbl> <dbl>
## 1 0.762 0.793 0.447
```

```
Genus_B %>%
  filter(Genus != "Unspecified") %>%
  summarise_if(is.numeric, sum, na.rm = TRUE)
```

```
## # A tibble: 1 x 3
##   `B-1` `B-2` `B-3`
##   <dbl> <dbl> <dbl>
## 1 0.526 0.533 0.547
```

```
Genus_C  %>%
  filter(Genus != "Unspecified") %>%
  summarise_if(is.numeric, sum, na.rm = TRUE)
```

```
## # A tibble: 1 x 3
##   `C-1` `C-2` `C-3`
##   <dbl> <dbl> <dbl>
## 1 0.920 0.972 0.972
```

```
Genus_D %>%
  filter(Genus != "Unspecified") %>%
  summarise_if(is.numeric, sum, na.rm = TRUE)
```

```
## # A tibble: 1 x 3
##   `D-1` `D-2` `D-3`
##   <dbl> <dbl> <dbl>
## 1  1.03 0.998 0.999
```

```
Genus_E  %>%
  filter(Genus != "Unspecified") %>%
  summarise_if(is.numeric, sum, na.rm = TRUE)
```

```
## # A tibble: 1 x 3
##   `E-1` `E-2` `E-3`
##   <dbl> <dbl> <dbl>
## 1 0.857 0.789 0.841
```

```
Genus_F %>%
  filter(Genus != "unclassified") %>%
  summarise_if(is.numeric, sum, na.rm = TRUE)
```

```
## # A tibble: 1 x 3
##   `F-1` `F-2` `F-3`
##   <dbl> <dbl> <dbl>
## 1     1  1.00     1
```

```
Genus_G  %>%
  filter(Genus != "Unspecified") %>%
  summarise_if(is.numeric, sum, na.rm = TRUE)
```

```
## # A tibble: 1 x 2
##   `G-1` `G-2`
##   <dbl> <dbl>
## 1 0.941 0.938
```

```
NIST_16SData  %>%
  filter(Genus != "unclassified") %>%
  summarise_if(is.numeric, sum, na.rm = TRUE)
```

```
## # A tibble: 4 x 4
##   Sample  Count    Reads RelAbund
##   <chr>   <dbl>    <dbl>    <dbl>
## 1 NIST-1 180835 24037500    0.940
## 2 NIST-2 145511 19511875    0.932
## 3 NIST-3 159003 21304625    0.933
## 4 NIST-4 222080 29659500    0.936
```

```
RA_NIST_SGData %>%
  filter(Genus != "unclassified") %>%
  summarise_if(is.numeric, sum, na.rm = TRUE)
```

```
## # A tibble: 20 x 2
##    Sample RelAbund
##    <chr>     <dbl>
##  1 1-1        1.00
##  2 1-2        1.00
##  3 2-1        1.00
##  4 2-2        1.00
##  5 3-1        1.00
##  6 3-2        1.00
##  7 4-1        1.00
##  8 4-2        1   
##  9 5-1        1.00
## 10 5-2        1.00
## 11 6-1        1.00
## 12 6-2        1.00
## 13 7-1        1.00
## 14 7-2        1.00
## 15 8-1        1.00
## 16 8-2        1.00
## 17 NIST-5     1.00
## 18 NIST-6     1.00
## 19 NIST-7     1.00
## 20 NIST-8     1.00
```

### genera uniquely identified in sample A-1, semi join shows that atleast one replicate (from not company A) also identified these Genera

```
CompanyA_outlier_comparison <- Donor5.Genus.Table %>%
  gather(key = Genus, value = "RelAbund", -Sample) %>%
  spread(key = Sample, value = "RelAbund") %>% 
  select( - "A-3_DTC") %>%
  semi_join(Genus.A3, ., by = "Genus")

anti_join(Genus.A3,CompanyA_outlier_comparison, by = "Genus")
```

```
## # A tibble: 0 x 2
## # … with 2 variables: Genus <chr>, A-3 <dbl>
```

```
save.image("NIST-analysis.RData")
```

```
save.image("NIST-analysis.RData")
```
